# Recycling of Bacterial RNA Polymerase by the Swi2/Snf2 ATPase RapA

**DOI:** 10.1101/2023.03.22.533849

**Authors:** Koe Inlow, Debora Tenenbaum, Larry J. Friedman, Jane Kondev, Jeff Gelles

## Abstract

Free-living bacteria have regulatory systems that can quickly reprogram gene transcription in response to changes in cellular environment. The RapA ATPase, a prokaryotic homolog of the eukaryote Swi2/Snf2 chromatin remodeling complex, may facilitate such reprogramming, but the mechanisms by which it does so is unclear. We used multi-wavelength single-molecule fluorescence microscopy in vitro to examine RapA function in the *E. coli* transcription cycle. In our experiments, RapA at < 5 nM concentration did not appear to alter transcription initiation, elongation, or intrinsic termination. Instead, we directly observed a single RapA molecule bind specifically to the kinetically stable post-termination complex (PTC) -- consisting of core RNA polymerase (RNAP) bound to dsDNA -- and efficiently remove RNAP from DNA within seconds in an ATP-hydrolysis-dependent reaction. Kinetic analysis elucidates the process through which RapA locates the PTC and the key mechanistic intermediates that bind and hydrolyze ATP. This study defines how RapA participates in the transcription cycle between termination and initiation and suggests that RapA helps set the balance between global RNAP recycling and local transcription re-initiation in proteobacterial genomes.

**SIGNIFICANCE:** RNA synthesis is an essential conduit of genetic information in all organisms. After transcribing an RNA, the bacterial RNA polymerase (RNAP) must be reused to make subsequent RNAs, but the steps that enable RNAP reuse are unclear. We directly observed the dynamics of individual molecules of fluorescently labeled RNAP and the enzyme RapA as they colocalized with DNA during and after RNA synthesis. Our studies show that RapA uses ATP hydrolysis to remove RNAP from DNA after the RNA is released from RNAP and reveal essential features of the mechanism by which this removal occurs. These studies fill in key missing pieces in our current understanding of the events that occur after RNA is released and that enable RNAP reuse.

## INTRODUCTION

DNA transcription into RNA by RNA polymerases (RNAP) is essential for protein synthesis and is universal in all domains of life. In *E. coli,* transcription is regulated at each step to control initiation (1), elongation (2, 3), and importantly, efficiency of termination (4, 5). Intrinsic termination occurs at sites on DNA which typically encode a hairpin structure followed by a short uridine-rich sequence. The formation of this RNA hairpin in the RNAP exit channel causes spontaneous transcript release as a result of melting and shearing of the RNA-DNA hybrid (6–9). The canonical scheme depicts transcript release followed by RNAP dissociation from DNA as the final step in the transcription cycle (5, 8), with RNAP free to bind a σ factor and initiate another round of transcription. RNAP recycling after termination is essential to maintain an adequate rate of steady-state transcription in vivo.

Contrary to the canonical scheme, recent single-molecule studies on the behavior of RNAP at intrinsic terminators demonstrate the formation of a long-lived RNAP-DNA complex, the post-termination complex (PTC), that persists on DNA after transcript release in >90% of transcription events in vitro (10–13). Consistent with these observations, ensemble experiments show that the PTC must undergo further conformational changes to drive RNAP dissociation from DNA (14). While in the PTC state, RNAP may hop or slide along DNA in a random walk and flip 180 degrees on DNA, allowing re-initiation of either sense or anti-sense transcription from a suitably situated promoter upon σ factor binding (10, 12, 13). However, these re-initiation pathways are limited to 1-D diffusion to proximal promoters on the same template, and efficiency of re-initiation through this pathway depends on terminator-promoter distance and σ factor availability (15). The mechanism by which RNAP dissociates from DNA during intrinsic termination and is efficiently recycled to the cytoplasmic pool of RNAP remains unknown.

RapA, <u>R</u>NAP-<u>a</u>ssociated <u>p</u>rotein <u>A</u>TPase, is a 109 kDa protein that is present across eubacteria and is as abundant in the cell as the housekeeping sigma factor σ^70^ (16). RapA stably associates with core RNAP (*K*D ∼5-10 nM) (16), but not with holoenzyme (σ^70^RNAP) due to the overlapping binding regions of RapA and σ^70^ on RNAP (17). The ATPase activity of RapA is significantly enhanced upon binding to RNAP compared to its apo (unbound) form (18), but is unaffected by presence of DNA. RapA is a bacterial homolog of eukaryotic Swi2/Snf2 proteins, which bind to and remodel nucleosomes on DNA (19). However, unlike some Snf2 family members, RapA binds dsDNA only weakly (*K*D ∼22 μM) (20, 21).

In bulk transcription reactions with purified RNAP and σ^70^, the addition of RapA strongly stimulates multi-round transcription cycling and transcript production, raising the possibility that it acts following termination (17, 22–25). However, multiple other activities for RapA have been proposed, including stabilizing open-promoter complexes during transcription initiation (26), promoting nascent transcript release from elongation complexes (24), preventing binding of RNAP on non-promoter DNA (27), and removing stalled elongation complexes via backtracking (28).

To directly examine the role of RapA in transcription in real time, we performed colocalization single-molecule spectroscopy (CoSMoS) (29, 30) and single-molecule fluorescence resonance energy transfer (FRET) (31) studies. These experiments allowed us to define the binding target of RapA in the transcription cycle, establish how RapA reaches this target, observe the formation of key RapA•RNAP•DNA complexes, and quantitatively characterize the kinetic pathways by which they form and resolve and the roles of nucleotide binding and hydrolysis. Together, the studies define a molecular mechanism for RapA function in bacterial transcription.

## RESULTS

### RapA binds to the PTC and promotes PTC dissociation

To observe the behavior and function of RapA during the transcription cycle by single- molecule fluorescence, we first tethered fluorescently labeled circular DNA molecules (DNA^Cy5^) containing a transcription promoter and two consecutive terminators to the surface of a glass flow chamber (Fig. 1A). Locations of the DNA^Cy5^ molecules were recorded by single-molecule TIRF microscopy, and the attached Cy5 dye moieties were then photobleached (see Methods). We then added to the chamber a solution containing σ^70^RNAP holoenzyme fluorescently labeled on the RNAP β′ subunit (σ^70^RNAP^549^). We observed the appearance of discrete RNAP fluorescent spots colocalized at DNA^Cy5^ locations, indicating the formation of open promoter complexes on DNA (11). Excess unbound RNAP was then flushed from the chamber.

**Figure 1.**
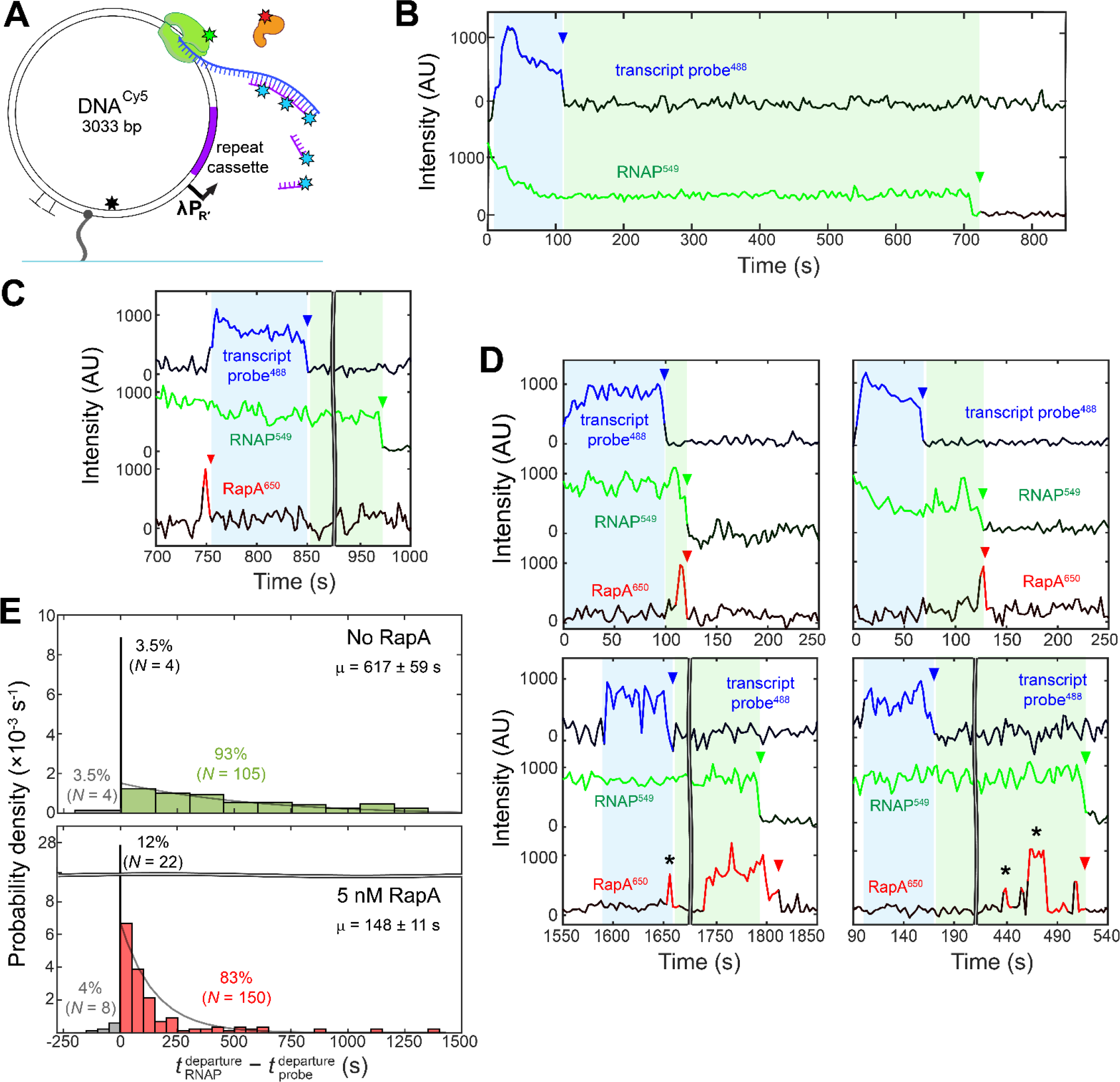
RapA removes post-termination RNAP from DNA. (A) Schematic of the single-molecule transcription experiment. Circular DNA^Cy5^ molecules tethered to a glass surface (blue) contain the λPR′ promoter, seven tandem repeats encoding a 21-nt hybridization target (purple), and 2 consecutive intrinsic transcription terminators (TT). DNA^Cy5^ molecules were photobleached (black star) after recording their locations. The presence at the DNA location of σ^70^RNAP^549^ (green), RapA^650^ (orange) or nascent transcript hybridization probe^488^ (blue) is detected by the fluorescence from the attached green-, red-, and blue-excited dye molecules (stars) under alternating blue and green+red laser excitation. Time resolution was 2.6 s. **(B)** Control transcription experiment without RapA. Plot shows an example two-color record of fluorescence intensity at the location of a single DNA molecule. Presence (color) or absence (black) of a DNA- colocalized fluorescence spot from the specified molecule is indicated. Simultaneous presence of RNAP^549^ and probe^488^ (blue shaded interval) reports presence of a transcription elongation complex. Departure of probe^488^ (blue arrow) indicates transcript release and thus termination. RNAP^549^ can persist on DNA for extended times after termination, before its eventual dissociation. Time interval (green shading) between probe^488^ departure and RNAP^549^ departure (green arrow) reports the presence of the post-termination complex (PTC). **(C, D)** Same as (B), but from a transcription experiment that includes 5 nM RapA^650^. Examples are shown of three-color intensity records at five DNA locations at which RapA^650^ binds and quickly departs (red arrow) during initiation or early elongation (C) or binds shortly before and departs simultaneously with or just after RNAP^549^ leaves the PTC (D). Asterisks mark non-productive RapA departures that did not drive PTC dissociation. **(E)** Normalized histograms of PTC lifetimes (green shaded intervals from (B, C, D)) in transcription experiments with zero (top) or 5 nM RapA (bottom). RNAP^549^ departed either before (grey), simultaneously (± 1 s) with (black), or after (green/pink) probe^488^ departure. Curves are exponential fits to lifetimes > 1 s yielding the indicated time constants μ. For the zero RapA experiment, the reported time constant underestimates the mean PTC lifetime because 1) for a minority of the observed probe departures no RNAP departure was observed before the end of the experimental record (26 of 113 total; these are not shown and were not included in the fit), and 2) the measured RNAP^549^ photobleaching rate of 2.4 ± 0.2 × 10^-4^ s^-1^(Fig. S4), reduces the measured characteristic lifetime by ∼15%.

We first did a control experiment without RapA in which we initiated transcription at time zero upon the addition of a solution containing 0.5 mM each of ATP, CTP, GTP, and UTP (NTPs), and 10 nM of an oligonucleotide probe labelled with Alexa Fluor 488 (probe^488^) which hybridizes to a repeat cassette sequence near the 5′-end of the nascent transcript. At a typical DNA location that started with bound RNAP^549^ (e.g., Fig. 1B) we observed fluorescence resulting from probe^488^ hybridization to the nascent RNA, indicating the existence of a transcript elongation complex (11, 32, 33). During the time that probe fluorescence was present, we sometimes observed that RNAP^549^ intensity gradually decreased as observed previously for transcription with linear DNA templates (11). We attribute this change to gradual movement of the RNAP^549^ away from the chamber surface in the direction of decreasing TIRF field intensity. On other DNAs we saw little intensity change during transcription, suggesting that some DNA circles might be oriented largely along the surface rather than normal to it. In either case, we usually observed the all-at-once disappearance of the probe fluorescence (Fig. 1B, blue arrow) suggesting that transcription had terminated via release of the probe-bound RNA. The mean duration of probe^AF488^ fluorescence was 59 ± 7 s (± SE; *N* = 113) for our experiment conducted at 33.1 °C, which is similar to the time (∼42-60 s) for *E. coli* RNAP to transcribe the 2,334 bp DNA estimated from literature elongation rates (39-55 nt s^-1^ at 37 °C (34–36)). After termination, RNAP typically persisted on the DNA for hundreds of seconds as a PTC.

To examine the role of RapA, we prepared the fluorescently labeled construct RapA^650^. When we added 5 nM RapA^650^ and 1 mM ATP to a chamber containing only DNA with no RNAP, we observed little RapA binding to DNA; 4% (9 of 226) of DNA molecules were observed to interact with RapA for at least 2 consecutive frames (2 s) over 500 s. In contrast, in a transcription experiment with 5 nM RapA^650^ (Fig. 1A) 66% (184 of 280) of the DNA locations exhibited RapA binding, suggesting a direct interaction of RapA with polymerase molecules bound to the DNA. Furthermore, RapA binding at DNA locations depended on the phase of the transcription cycle. For the 180 of the 280 DNA molecules that exhibited transcript production, (i.e., probe appearance and later disappearance) we observed in 55 cases (31%) a brief RapA association just prior to probe appearance (e.g., Fig. 1C). The significance of this short-lived binding during initiation or early elongation is not known (see Discussion). Importantly, RapA rarely bound to mature elongation complexes. RapA presence during intervals when probe was colocalized, which requires a transcript length of > 47 nucleotides, was seen on only 10% (18 of 180) of DNAs that exhibited probe binding. Furthermore, the average dwell time of RapA on the elongation complexes to which it did bind was brief, 13 ± 3 s (± SEM; *N* = 18). Consistent with these observations, the presence of 5 nM RapA^650^ had little effect on the rate of transcript elongation: the mean duration of the probe^AF488^ fluorescence was 68 ± 3 s (± SEM; *N* = 180), similar to that seen in the absence of RapA.

In contrast to the minimal binding observed during transcript elongation, most (76 %; 136 of 180) transcription events displayed RapA binding that occurred simultaneously with (within the ± 2.6 s time resolution of the experiment) or after transcription termination detected by probe^AF488^ departure (examples shown in Fig. 1D). Notably, the majority of RNAP releases were preceded by RapA binding and occurred simultaneously with RapA departure from DNA (60%; 107 of 180 transcription events). Occasionally, RapA^650^ stayed bound on DNA after RNAP^549^ departure (Fig 1D, bottom left), but these events were rare (4%; 7 of 180) and may be due to RNAP^549^ photobleaching. Thus, RapA appears to bind to the PTC and remove RNAP from the DNA. This effect is confirmed by measurements of PTC lifetimes. Without RapA, PTCs had a characteristic lifetime of > 600 s (Fig. 1E, top), similar to that observed previously (10). These measurements of PTC lifetimes without RapA are underestimates of the true mean because 23% (26 of 113 transcription events) of RNAP molecules stayed bound until the end of the experimental record. Not all RapA binding events triggered RNAP release; many were non-productive events that ended with RapA departure from DNA with RNAP remaining behind (e.g., Fig. 1D, stars). Nonetheless, inclusion of 5 nM RapA^650^ in the experiment decreased PTC lifetimes by more than four-fold (Fig. 1E, bottom). In summary, the data establish via direct observation of transcription complexes that low nanomolar concentrations of RapA preferentially bind the PTC, are present when core departs, and always or almost always leave with the departing RNAP.

### PTC lifetimes are RapA concentration- and ATP-dependent

To further investigate the mechanism of PTC dissociation, we next reconstituted PTCs. The PTC consists of core RNAP bound sequence-nonspecifically to DNA. Available data suggest that the same complex can be made directly by mixing core RNAP directly with DNA. Like PTCs, these reconstituted PTCs (rPTCs) have long lifetimes (>600 s), exhibit RNAP sliding on DNA, are dissociated by treatment with heparin, and can bind σ^70^ (10, 11, 15). rPTCs can be efficiently made in large quantities at the beginning of an experiment, allowing us to systematically investigate the effect of RapA concentrations and ATP on rPTC lifetimes.

We first tethered to a coverslip fluorescently labeled DNA circles that lack known transcription promoters, npDNA^Cy5^, and then preincubated with RNAP^549^ (Fig. 2A). Free RNAP was flushed from the chamber and replaced by a solution containing various concentrations of RapA and/or ATP at time zero. We then measured the dwell times of RNAP^549^ spots at npDNA^Cy5^ locations (Fig. 2B). At zero RapA and 1 mM ATP, the rPTC mean dwell time was 1,271 ± 26 s (SEM; N = 1063), but this underestimates the true mean since 41 % (442 of 1063) of measurements were censored by the end of the experimental record. This value is consistent with the > 600 s lifetime observed for PTCs (Fig. 1). When RapA in the 1 – 10 nM range was added the reciprocal mean dwell time increased steeply (Fig. 2C, green circles). In contrast, no change in mean dwell time was observed at increasing concentrations of RapA in the absence of ATP (Fig. 2C, green diamonds).

**Figure 2.**
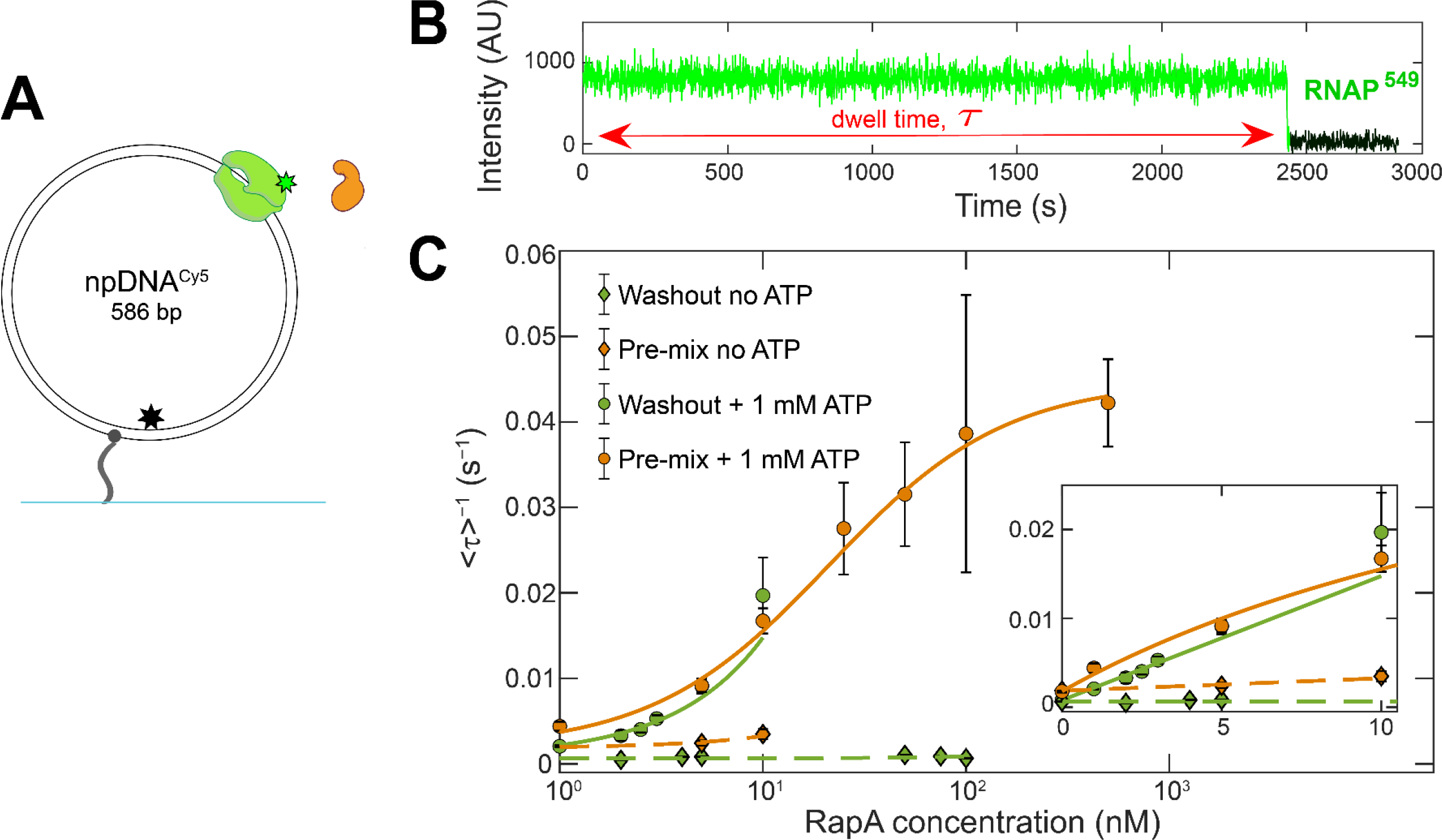
Nanomolar RapA disrupts PTCs in an ATP-dependent manner. (A) Schematic of the single- molecule rPTC disruption experiment. Circular npDNA^Cy5^ molecules (tethered to a glass surface (blue)) contain no known promoter sequences and were photobleached (black star) after recording their locations. At time zero, RapA or RapA^650^ (orange) was added either together with RNAP^549^ (green; pre-mix) or after free RNAP^549^ was removed from the chamber (washout). Presence at the DNA location of RNAP^549^ is detected by the fluorescence from the green-excited dye molecule (star) under continuous laser excitation. Time resolution 1 s; 532 nm laser power 400 µW; RNAP^549^ photobleaching rate (Fig. S3) 0.0006 s^-1^. **(B)** Example RNAP^549^ fluorescence intensity record at the location of a single DNA molecule from a washout experiment, plotted as in Fig. 1B-D. **(C)** Reciprocal of the mean RNAP^549^ dwell time τ at various RapA or RapA^650^ concentrations. Experiments were conducted using the pre-mix or washout protocols in the presence or absence of 1 mM ATP. Data were fit to hyperbolic (solid line) or linear (dashed lines) models yielding fit parameters given in Table S1. Inset: Magnified view of the low concentration range with additional data at zero RapA.

PTCs dwell times were the same with 5 nM unlabeled RapA and 5 nM RapA^650^ (Fig. S1), indicating that the presence of the fluorescent dye label on RapA does not alter its PTC disruption activity.

At RapA concentrations > 10 nM in the presence of ATP, the dwell time of RNAP on DNA became sufficiently short that significant dissociation occurred during the washout fluid flow. To measure short dwell times accurately under these conditions, we used an alternative “pre-mix” protocol in which 0.7 nM RNAP^549^ and various concentrations of RapA or RapA^650^ were pre- mixed with or without ATP, and then loaded all together over the surface-tethered npDNA molecules. RNAP^549^ molecules were continuously recorded as they bound and released from DNA surface locations, allowing measurements of dwell times as short as 2 s. When the two protocols were compared under the same conditions, they produced similar results (Fig. 2C, orange and green). The dwell times measured in the pre-mix experiment exhibited saturation kinetics and fit well to a hyperbolic model with a half-saturating RapA concentration *K*0.5 = 21 ± 3 nM and extrapolated maximum rate at saturation of *k*max = 0.043 ± 0.004 s^-1^ (Table S1). These parameters imply an effective second-order rate constant for RapA-mediated rPTC disruption of *k*max / *K*0.5 = (2.1 ± 1.3) × 10^6^ M^-1^ s^-1^, close to that expected for a diffusion-limited association step. Taken together, these data indicate that nanomolar concentrations of RapA potently disrupt rPTCs in an ATP-dependent manner; RapA is able to decrease PTC lifetime by a factor of at least 20-fold (*k*max / *b*; see Table S1).

Washout experiments in which we observed fluorescence from both RNAP^549^ and RapA^650^ (Fig. 3A) exhibited similar behavior on rPTCs to that seen on PTCs (Fig. 1D). RapA bound to rPTCs and usually departed simultaneously with RNAP. In rare cases, RapA^650^ remained briefly after RNAP^549^ fluorescence vanished, but as previously noted these uncommon events may be a consequence of RNAP^549^ photobleaching rather than true sequential departure of the two proteins. With both PTCs (Fig. 1D, asterisks) and rPTCs (Fig. 3A, asterisks), we sometimes observed non-productive RapA binding that did not remove RNAP from DNA.

**Figure 3.**
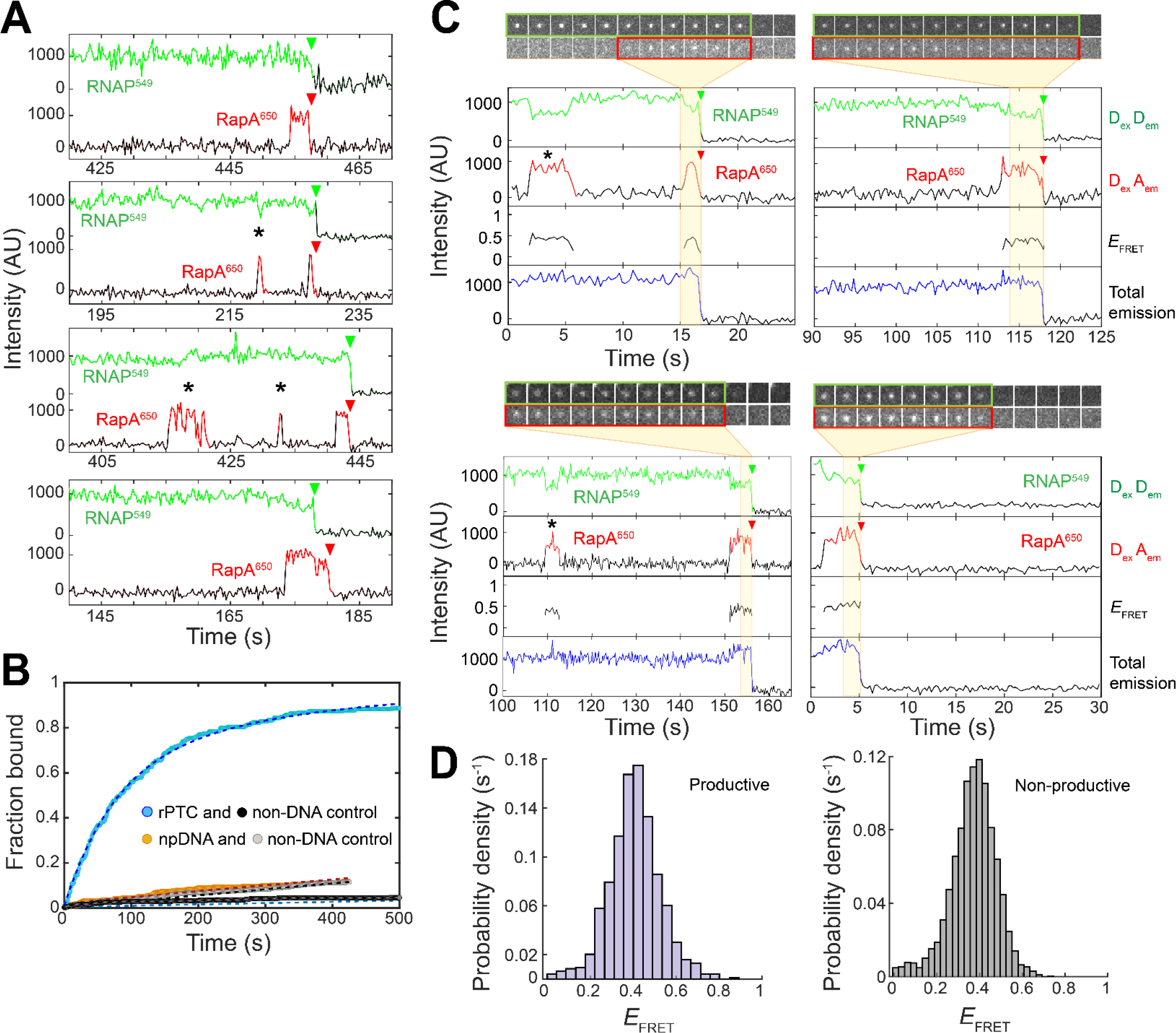
Dynamics of labeled RapA interacting with rPTCs. (A) Example fluorescence intensity records from the locations of four individual rPTCs in the presence of 0.75 nM RNAP^549^, 5 nM RapA^650^, and 1 mM ATP. Experiments used the design shown in Fig. 2A with the washout protocol, and data were collected with simultaneous green and red laser excitation; time resolution 0.2 s. Asterisks and arrowheads respectively indicate non-productive binding events by RapA and protein departure, respectively, as in Fig. 1. **(B)** Cumulative fraction of rPTC locations from the experiment in (A) that bound at least one RapA^650^ molecule by the indicated time (blue) and analogous data from non-DNA control locations in the same experiment. Plot truncated at 500s for clarity. Also shown are data from an analogous experiment without RNAP that show binding to npDNA locations (orange) and non-DNA control locations. Fits to model that includes exponential non-specific surface binding and exponential DNA-specific binding (dashed lines) yielded the kinetic parameters shown in Table S2. **(C)** Experiment with same conditions as in (A), except that data were recorded with only 532 nm laser excitation. Example records from four different PTC locations show donor emission with donor excitation (DexDem), acceptor emission with donor excitation (DexAem), effective FRET efficiency (*E*FRET = DexAem / (DexDem + DexAem)), and total emission (DexDem + DexAem). Galleries show excerpts of the PTC location image recordings in the DexDem and DexAem channels. **(D)** Histograms of *E*FRET values from the experiment in (C) at times when both RNAP^549^ and RapA^650^ were detected on PTCs. Shown separately are data from RapA dwells that ended with simultaneous RapA and RNAP dissociation (left, mean *E*FRET = 0.404 ± 0.004 (SEM), *N* = 932 time points from 46 PTCs) and those that ended with nonproductive RapA dissociation (right, mean *E*FRET = 0.370 ± 0.002, N = 4,253 time points from 94 complexes).

### RapA binds directly to core RNAP on DNA

We next investigated the steps in the process by which RapA disrupts PTCs. Since RapA has putative DNA-binding domains (17, 18), we first asked whether the protein initially binds DNA or alternatively whether it goes directly to the DNA-bound RNAP in the PTC. In the Fig. 3A experiment, RapA^650^ bound to rPTCs with a second-order rate constant of 1.8 ± 0.4 × 10^6^ M^-1^ s^-1^ (Fig. 3B; Table S2). This is similar to the effective second order rate constant for PTC disruption in the limit of low RapA concentrations (*k*max / *K*M = 2.1 ± 1.3 × 10^6^ M^-1^ s^-1^; Fig. 2C and Table S1), confirming that the observed binding is kinetically competent to be the initial step in PTC disruption.

In contrast to its rapid binding to PTCs, 5 nM RapA^650^ in 1 mM ATP exhibited no detectable background binding to the slide surface or to DNA molecules without bound RNAP (Fig. 3B, compare orange and gray distributions; Table S2, compare *A*f values). This is consistent with the very weak binding of dsDNA by RapA in solution (*K*D = 15 µM (18)). Thus, RapA likely reaches its PTC target by direct binding from solution, not by long-distance random sliding on DNA until RNAP is encountered.

To verify that RapA and RNAP are in close proximity when RapA binds to rPTCs, we checked for FRET between RNAP^549^ (donor) and RapA^650^ (acceptor). When washout protocol experiments were repeated using only donor (532 nm) excitation, we observed appearance of DNA-colocalized spots in the acceptor emission channel and concomitant reduction of donor emission, both indicative of FRET (Fig. 3C). The effective FRET efficiency (*E*FRET) distribution was unimodal (Fig. 3D, left), with mean <*E*FRET> = ∼0.4, a plausible value given the proximity of the SNAP-tag attachment points in the two labeled proteins in RapA-elongation complex structures (Fig. S2).

Just as we observed for PTCs (Fig. 1D), binding of RapA to rPTCs was sometimes productive, ending with simultaneous release of RapA and RNAP, and sometimes non-productive, ending with just RapA release (Figs. 3A and 3C, red arrows and asterisks, respectively). *E*FRET distributions for the productive and non-productive RapA•RNAP•DNA complexes were similar (Fig. 3D), consistent with the idea that productive and non-productive events reflect different breakdown pathways of the same types of RapA-rPTC complexes.

### RapA association with PTC is independent of nucleotide

Many members of the Snf2 family are motor enzymes that use ATP hydrolysis to drive translocation of the Snf2 homology domain relative to DNA. To check if RapA translocates along DNA to reach DNA-bound RNAP, we compared RapA recruitment to PTCs in experiments with ATP, with no added nucleotide, or with the non-/slowly-hydrolysable ATP analog AMPPNP (Fig. 4A). For each rPTC present at the beginning of the washout experiment (e.g., as seen in Fig. 4A, DexDem records), we measured the interval up to the time that RapA associated with the PTC as judged by the onset of FRET (DexAem). Fitting demonstrates that the PTC-specific binding intervals are exponentially distributed (Fig. 4B), as expected for a simple, one-step binding process. Importantly, the RapA-PTC association rate constants *k*1 determined from the fits were identical within experimental uncertainty (Fig. 4C). Thus, the absence of nucleotide or nucleotide hydrolysis does not alter the kinetics of RapA association with the PTC, implying that translocation driven by RapA ATP hydrolysis is not required for RapA recruitment to the PTC. Furthermore, the onset of RapA^650^ binding detected by direct excitation (Fig. 4A, AexAem) was simultaneous with the onset of the same binding event detected by FRET (DexAem). The immediate appearance of FRET simultaneously with binding is consistent with direct association of RapA with DNA-bound RNAP. The lack of a detectable delay between RapA arrival and FRET shows either that there is no intermediate state in which RapA searches along DNA, or that this intermediate is too short-lived to detect at the ∼1.4 s time resolution of the experiment.

**Figure 4.**
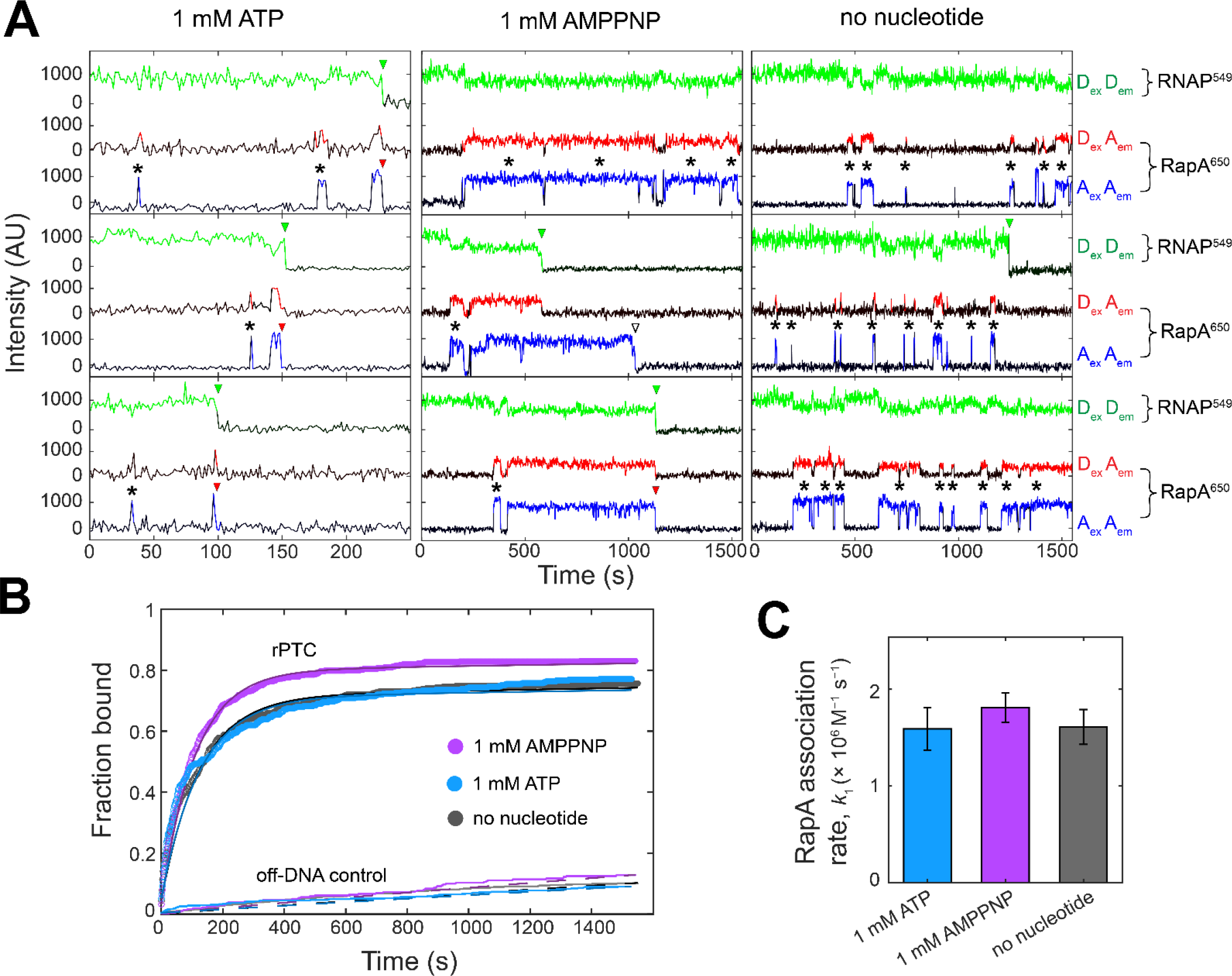
RapA-PTC association kinetics under different nucleotide conditions. (A) Example fluorescence intensity records from nine individual rPTC locations in washout experiments (as in Fig. 2A, B) with either 1 mM ATP (left), 1 mM AMPPNP (center), or no added nucleotide (right). Alternating excitation with green and red lasers (0.2 s duration frames) allowed detection of RNAP^549^ (DexDem), RapA^650^ excited via FRET (DexAem), and RapA^650^ excited directly (AexAem). The average interval between consecutive frames with the same excitation wavelength was 1.39 ± 0.03 (SD) s. Note the expanded horizontal scale for the 1 mM ATP condition. Departure or photobleaching of core RNAP^549^ is marked by green arrows; departure of RapA^650^ without simultaneous RNAP^549^ release in marked by an open arrow. Asterisks mark non-productive RapA binding intervals (RapA binding without catalyzing an RNAP departure). **(B)** Kinetics of specific RapA-rPTC association detected by FRET (i.e., DexAem). For the same three experiments from which the examples in (A) were taken, plots show the cumulative fraction of rPTCs that exhibited at least one RapA^650^ binding by the indicated time (rPTC) and analogous data from control locations without DNA molecules in the same samples (off-DNA control). Fits to data from both the rPTC (solid lines) and off-DNA data (dashed lines) using the same model as in Fig. 3B yield the parameters in Table S3. **(C)** Second-order association rate constants for RapA binding to PTCs (see Table S3).

### Two different RapA•RNAP•DNA complexes form during PTC disruption

The forgoing experiments imply that the mechanism of PTC disruption by RapA starts with RapA binding to the RNAP component of the PTC and ends with simultaneous departure of RapA and RNAP from the DNA. We next examined the characteristics of the RapA•RNAP•DNA complexes that exist in the time interval between these two events. In contrast to the similar rates of RapA•RNAP•DNA complex formation in different nucleotide conditions, RapA•RNAP•DNA lifetimes were markedly different. Notably, 1 mM AMPPNP caused a >50-fold increase in the average RapA•RNAP•DNA lifetime relative to ATP (373 ± 22 (SEM) s, *N* = 645 and 7.1 ± 0.4 s, *N* = 647, respectively). Under all three nucleotide conditions, individual RapA•RNAP•DNA complexes were observed to have a broad distribution of lifetimes. Each distribution had two (or more) exponential components (Fig. 5A). This demonstrates that during the PTC disruption reaction, at least two different RapA•RNAP•DNA complexes are formed.

**Figure 5.**
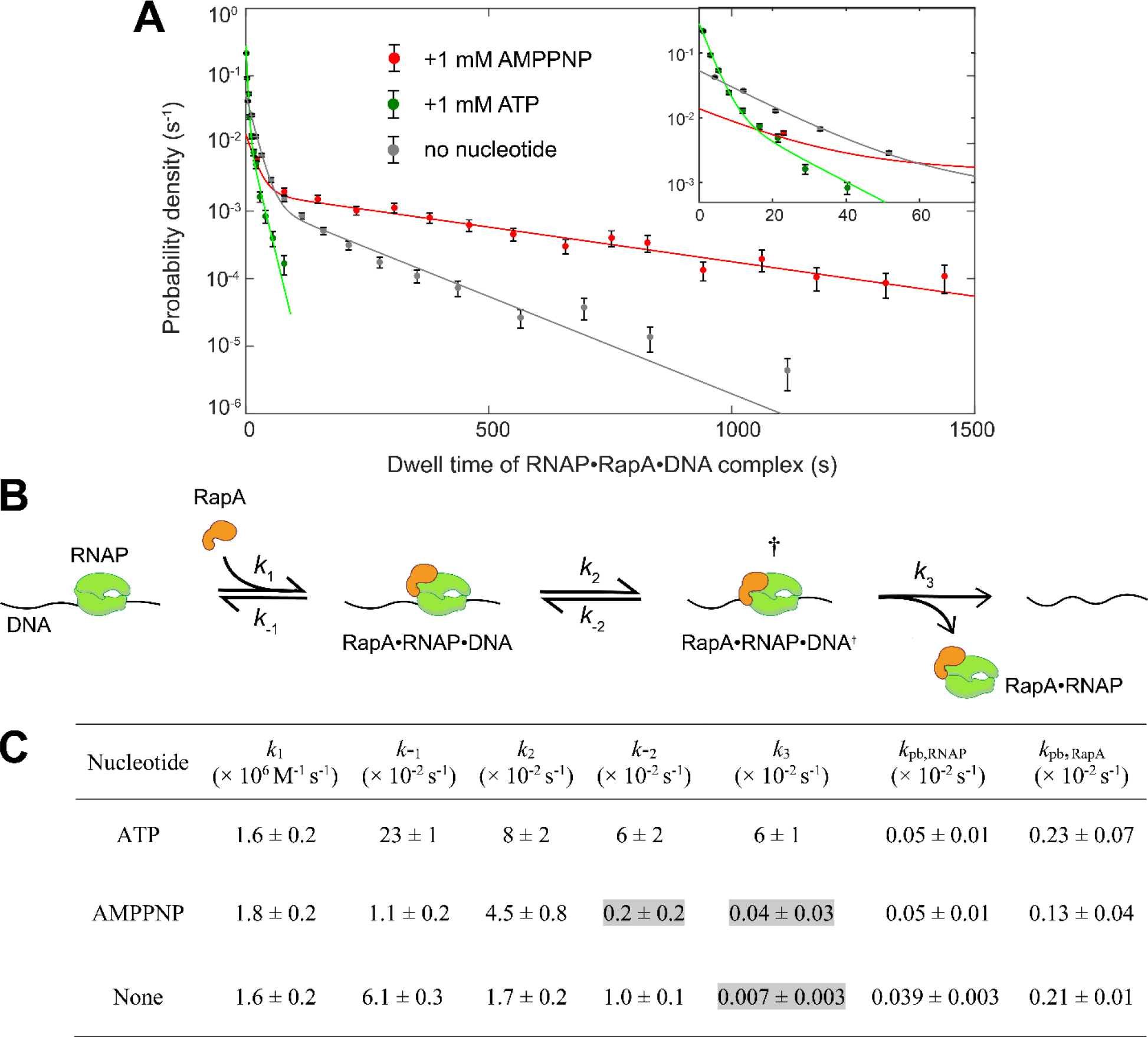
Kinetic analysis of PTC disruption by RapA. (A) Lifetime distributions of all RapA^650^•RNAP^549^•DNA complexes (circles; ± S.E.) for the different nucleotide condition experiments shown in Fig. 4. Also shown (lines) are maximum-likelihood fits to a two-exponential equation (see Methods); fit parameters are provided in Table S7. **(B)** Summary proposed kinetic scheme for RNAP (green) dissociation from the PTC by RapA (orange). The RapA•RNAP•DNA and RapA•RNAP•DNA^†^ complexes are indistinguishable by fluorescence, but the presence of two such species is inferred from the kinetics (see text). Experimental data were fit with a full scheme that includes additional steps accounting for photobleaching of RapA^650^ and RNAP^549^ (Fig. S5). In our working model of the mechanism, ATP binds during the RapA•RNAP•DNA state and is cleaved upon entrance into or exit from the RapA•RNAP•DNA^†^ state (see Discussion). **(C)** Rate constants for the steps in the scheme (B) and for the photobleaching steps (see Fig. S5). Rate constants were determined by fitting the full kinetic scheme (Fig. S5) to the RapA•RNAP•DNA paired lifetime and fate data summarized in Fig. S6 and Table S6, except for *k*1 values which are from Table S3. Gray highlights designate values that are indistinguishable from zero within experimental uncertainty. The given photobleaching rate constants were determined from the fits, but they are roughly similar to photobleaching rate constants measured independently: *k′*pb,RNAP = (0.043 ± 0.003) × 10^-2^ s^-1^ (Table S4) and *k′*pb,RapA = (0.26 ± 0.09) × 10^-2^ s^-1^ (Table S5). The lifetime distributions and fate partition ratios predicted by the fits to the model are shown in Fig. S6 and Table S6.

### Kinetic mechanism of PTC disruption by RapA

We propose (Fig 5B) a minimal kinetic scheme consistent with our observations, one in which the two ternary complexes (designated RapA•RNAP•DNA and RapA•RNAP•DNA^†^) are sequential intermediates in PTC disruption. We speculate that the two intermediates may differ in conformation, nucleotide state of the active site, or both. To test the agreement of this scheme with the experimental data, we first classified by its fate and measured its lifetime each individual observed RapA•RNAP•DNA complex. Three fates were observed: 1) the unproductive fate, in which RapA departs by itself (e.g., the RapA•RNAP•DNA complexes marked “*” in Fig. 4A); 2) the productive fate where RapA and RNAP leave simultaneously (e.g., Fig. 4A, red and green arrowheads); and 3) the fate in which RNAP departure precedes RapA departure (e.g., Fig. 4A, open arrowhead). We then fit the model and determined the values of the rate constants that gave the best agreement with the lifetime and fate data independently for each of the three experiments in Fig. 4A.

The model accurately reproduced the molecular behaviors observed for the RapA•RNAP•DNA complexes. In particular, it reproduced the partitioning between the three fates (Table S6) and the lifetime distributions for RapA•RNAP•DNA complexes segregated according to fate (Fig. S6). In our model, we considered the possibility that disappearance of the RapA^650^ or RNAP^549^ fluorescent spots can be caused either by protein dissociation or by photobleaching (Figure S5; see Methods). For RapA, we explicitly modeled dissociation (*k*-1) and photobleaching (*k*pb,RapA) processes. For RNAP we found that it was not necessary to include a process in which RNAP dissociates from a RapA•RNAP•DNA complex leaving RapA behind to give good agreement with the data, so such a step was omitted from the kinetic scheme (Fig. 5B; Fig. S5) The validity of the photobleaching analysis is confirmed by independent measurements of the photobleaching rates of RNAP^549^ (Fig. S3, Table S4) and RapA^650^ (Fig. S4, Table S5) which agree with the corresponding rate constants determined by fitting the RapA•RNAP•DNA lifetime and fate data to the kinetic scheme (Fig 5C).

The kinetics revealed in the quantitative analysis of the reaction scheme (Fig. 5C) emphasize the kinetic significance of the two RapA•RNAP•DNA complexes: both *k*2 and *k*3 are partially rate limiting for RapA function at 1 mM ATP and the reverse reactions *k*-1 and *k-*2 significantly compete with the *k*2 and *k3* forward processes. Thus, the RapA binding step and the subsequent isomerization step are readily reversible, and this accounts for the significant number of non-productive RapA binding events seen in the presence of ATP. In both AMPPNP and no nucleotide conditions, even more non-productive events are observed, although it is possible that some of these events are due to photochemical blinking of the dye on RapA^650^.

## DISCUSSION

Early work on RapA showed that it facilitated multi-round transcription in vitro, including switching between different template molecules. Jin and co-workers proposed that RapA allows recovery of RNAP from a “posttranscription or posttermination” refractory state, but the composition of this refractory state and its position in the transcription cycle remained unclear (17, 22, 23). Here we provide direct evidence that RapA binds to the post-termination RNAP-DNA complex (PTC), an intermediate in the transcription cycle that follows intrinsic termination (10, 12, 13, 15). Within seconds of its binding to the PTC, RapA effectively removes core RNAP from DNA. This process is driven by a single RapA molecule, occurs even at low nanomolar concentrations of RapA, is ATP-dependent, and does not require a sigma factor. The refractory state of the early studies seems similar to the PTC, but may not be identical to it. Notably, entrance into the refractory state appears largely irreversible, whereas PTCs can slowly dissociate even without RapA, especially in the presence of free σ^70^ (15).

By directly observing transcription events in real time, we were able to distinguish the step(s) during which RapA preferentially binds in the transcription cycle. We saw essentially no RapA binding to elongation complexes in our experimental conditions. This observation argues against suggestions that RapA acts directly on elongation complexes (28, 37, 38). Our finding is consistent with structures of the RapA·RNAP complex showing that RapA binds at the RNAP RNA exit channel and displaces the RNAP β-flap tip, thereby partially blocking the RNA exit channel. This likely makes RapA binding incompatible with the protruding RNA present in mature elongation complexes and also with the presence of elongation-complex-bound NusA (27, 37). Nevertheless, prior studies taken together with our results support an *indirect* RapA effect on elongation complexes. Portman et al. (28) saw RapA disrupt stalled elongation complexes, but primarily on positively supercoiled DNA, and only slowly (the minimal lifetime of elongation complexes with 100 nM RapA in solution was ∼1,000 s, not the 23 s we infer for PTCs at saturating RapA (1 / *k*max; see Table S1)). Importantly, they provided strong evidence that this activity of RapA requires elongation complex backsliding on RNA/DNA. Based on the structural data and our observations that RapA does not bind stably to mature elongation complexes, we propose that for an elongation complex to be disrupted, it must first backslide either off the 5′-end of the RNA or at least to within 8 nt from the 5′ end, the point at which the RNA-DNA heteroduplex starts to be disrupted. In either case, the result would be a PTC-like structure in which core RNAP is bound to DNA without RNA protruding from the exit channel and which is vulnerable to at least partial bubble-closing induced by positive superhelical twist. RapA could then bind the PTC-like structure and execute the ATP-dependent PTC-disruption activity that we demonstrate here. The fact that both elongation complex disruption and PTC disruption have half-maximal rates at similar RapA concentrations (∼ 7 nM) is consistent with this interpretation.

In addition to PTC binding, we also see RapA transiently (typically for < 4 s) associate with initiation complexes in ∼24% of transcription events. When RapA dissociates from these initiation complexes, the nascent RNA, and hence the elongation complex, is detected within seconds (Fig 1C). A similar function of RapA during initiation was also reported for *Pseudomonas putida* RapA, which is thought to stabilize the open complex during initiation and/or to rescue stalled initiation complexes, possibly in addition to its PTC disruption activity (39). Our data is consistent with a possible additional activity of *E. coli* RapA during initiation, although our experiments did not reveal an effect of nanomolar RapA on initiation kinetics.

The proposed kinetic scheme for disruption of the PTC by RapA (Fig. 5B) is a *minimal* scheme: it has the minimum number of states needed to explain our experimental results. The full mechanism for RapA function is necessarily more complex; in particular, it must also include ATP binding, hydrolysis, and ADP and Pi release steps. Our experiments conducted under various nucleotide conditions give some insight into the way that ATP hydrolysis is coupled to PTC disruption by RapA. The rate at which RapA associates with the PTC (*k*1 in Fig. 5B, C) is identical whether or not nucleotide is present, consistent with a mechanism in which RapA binds the PTC in its apo (no nucleotide) form, even when millimolar nucleotide is present. This is supported by structural work showing that the position of the RapA N-terminal domain (NTD) in apo-RapA may allosterically inhibit ATP binding (27, 38) and from experiments showing that deletion of RapA NTD significantly activates RapA ATPase activity in the absence of RNAP (40). Upon binding to RNAP either alone or in the context of an elongation complex, the NTD is repositioned, allosterically remodeling the RapA ATPase active site. This suggests that ATP binding to the active site occurs immediately after RapA binds to the PTC, consistent with our observation that all subsequent reactions of the RapA•RNAP•DNA complexes (*k*-1 and *k*2 in Fig. 5B, C in particular) have nucleotide dependent rates. Thus, ATP binding likely occurs between the *k*1 and *k*2 steps.

In contrast to the nucleotide-independent effect on *k*1, the nucleotide effect on *k*3 is profound. In the absence of nucleotide or in the presence of AMPPNP, *k*3 is not significantly above zero (Fig. 5C, gray). Based on the calculated uncertainties, we estimate that *k*3 is at least 150-fold faster in the presence of ATP than AMPPNP, and at least 900-fold faster than without nucleotide. To compare, the nucleotide effect on *k*2 is less than 5-fold. Therefore, we propose that the ATPase hydrolysis step is likely at or immediately prior to the step in which RapA-RNAP dissociates simultaneously from DNA, that is, that hydrolysis occurs upon entrance into or exit from the RapA•RNAP•DNA^†^ state shown in Fig. 5B.

RapA belongs to the Swi2/Snf2 protein family, many members of which are eukaryotic chromatin remodelers (19, 41). Some well-studied examples of these proteins function by a mechanism in which they bind to the histone core and use free energy from ATP hydrolysis to forcibly translocate or rotate the DNA, thereby disrupting histone-DNA contacts and facilitating displacement of the histones from DNA (41). Our data on RapA suggests that it works by an analogous mechanism in which it binds directly to the RNAP-DNA complex, followed by ATP binding and hydrolysis to drive RNAP ejection from DNA. Recent studies show the existence of a partially rewound DNA bubble in RNAP after RNA release during intrinsic termination (42). After binding to the PTC, RapA may mechanically close the bubble, thereby promoting PTC disruption. Further work will be required to determine if RapA perturbs or translocates on the DNA relative to RNAP during RapA•RNAP•DNA complex disruption, and if hydrolysis of only one molecule of ATP is sufficient for the process.

In bacterial transcription, there is evidence for re-initiation after intrinsic termination from experiments both in vitro and in vivo (10, 12, 13, 15). These studies, among others (43–46), are consistent with the proposal that there is a balance between re-initiation (local) and conventional initiation (global) in cells. Based on the activity and mechanism of RapA described here, we propose that the primary biological function of RapA is to set the balance between these two types of initiation. This phenomenon may be driven by RapA competition with sigma factors for binding to PTCs, since the binding sites of the two proteins on the β subunit of RNAP largely overlap (27, 37). Under particular growth conditions (e.g., log phase) transcription is more or less concentrated into particular genomic regions (e.g., rRNA operons) (43, 45, 47). Efficient re-initiation likely helps stabilize this localized transcription. However, when conditions change, it may confer a selective advantage for bacteria to be able to rapidly redeploy the transcription machinery to new genomic regions. In this context, it is noteworthy that RapA knockout mutants do not show any growth defects in unaltered rich medium. However, upon salt stress, sodium deoxycholate treatment, or osmotic shock, deletion of the *rapA* gene increased the duration of the recovery phase compared to WT (23, 48) and decreased antibiotic resistance in biofilms due to the lower expression of genes implicated in resistance (49). Furthermore, the RapA homolog HepA in *V. cholerae* is crucial for the acid-tolerance response, where *ΔhepA* mutants display a >1000-fold decrease in colonization proficiency (50). Consistent with RapA functioning in reprogramming, RapA is upregulated when cells transition into heat shock as modulated via the σ^32^ regulon (51). Further work will be needed to establish the role of RapA in transcriptional reprogramming.

## MATERIALS AND METHODS

### DNA and plasmids

Plasmid pDT2 (Addgene #199119) and pDT4 (Addgene #199120) are identical to pCDW114 and pCDW116 (10) respectively, except for mutation of 5′-CTGGAGTGCG-3′ to 5′-CTGGAGACCG-3′ to introduce a second BsaI site.

Circular DNAs were used in the single-molecule experiments to avoid adventitious binding of RNAP to the ends of linear dsDNAs (52). The circular DNA^Cy5^ templates used in transcription experiments were built using Golden Gate Assembly with pDT2 as the template and a synthetic duplex oligonucleotide containing internal biotin and Cy5 dye modifications. The 76 bp duplex was made by annealing 20 nM each of the complementary DNAs 5′-CGA TTA GGT CTC GGG CTA GTA CTG GTT TCT AGA G/iCy5/GT TCC AAG CC/iBio/ TCA CGG CGG CCG CCC ATC GAG ACC GGT TAA CC-3′ and 5′-GGT TAA CCG GTC TCG ATG GGC GGC CGC CGT GAG GCT TGG AAC CTC TAG AAA CCA GTA CTA GCC CGA GAC CTA ATC G-3′ (IDT). We performed a BsaI Golden Gate assembly (NEB, Golden Gate Assembly Mix) in T4 DNA Ligase Buffer in 20 µL volumes using equimolar pDT2 and the Cy5/biotin duplex at a final concentration of 6 nM each. Reactions were incubated for 35 alternating cycles of 5 min at 37 °C and 10 min at 16 °C, followed by 5 mins at 55 °C and then 10 min at 65 °C to inactivate T4 DNA ligase. To remove products that were not fully formed duplex closed circles, reactions were digested with 10 units of T5 exonuclease (NEB) for 30 min at 37 °C, followed by inactivation with 15 mM EDTA. The 3,033 bp product was further purified with Qiaquick PCR Purification Kit (Qiagen).

To make the 586 bp circular promoter-less template npDNA^Cy5^, we did the same as for DNA^Cy5^, except that the template used in Golden Gate Assembly was a PCR product amplified from pDT4 using primers 5′-GAA GGT CTC CAG CCG TAC CAA CCA GCG GCT TAT C-3′ and 5’-CCG GGT CTC ACC ATA CCC GCT GTC TGA GAT TAC G-3′.

Probe^488^, the 20-nt AlexaFluor-488 labelled transcription hybridization probe, was 5′-GTG TGT GGT CTG TGG TGT CT/3AlexF488N/-3′ (IDT).

### Proteins

The N-terminally His6-tagged *E.* coli RapA construct (H6-RapA) was expressed and purified as described (27).

From the H6-RapA expression plasmid and pSNAP-tag(T7) plasmid (NEB #101137), we constructed a plasmid pKI1 (Addgene #199118) encoding H6-SNAP-RapA. The plasmid was transformed into NEBExpress^IQ^ cells and the 129 kDa H6-SNAP-RapA protein was expressed and purified as described (27). The concentration of SNAP-RapA was calculated from absorbance using extinction coefficient, ε^280^ = 114,590 M^-1^ cm^-1^.

To make RapA^650^, the dye-labelled H6-SNAP-RapA construct, H6-SNAP-RapA protein and JFX- 650 fluorophore SNAP substrate (kind gift of Luke Lavis, Janelia Farm Research Campus) were mixed at a 1:2 molar ratio with 1 mM DTT in SNAP reaction buffer (50 mM Tris-HCl, pH 7.5, 100 mM NaCl, and 0.05% Tween-20). Excess dye was removed from solution using a Amicon Ultra-0.5 mL 30K spin column. The dye labelling efficiency was 80%, determined using the extinction coefficients of dye (ε^650^ = 160,000 M^-1^ cm^-1^ and ε^280^ / ε^650^ = 0.89) and protein (ε^280^ = 114,590 M^-1^ cm^-1^). RapA^650^ was frozen in liquid N2 and stored at -80°C in a buffer containing 10 mM Tris-HCl (pH 7.9), 50% glycerol, 0.1 mM EDTA, 0.1 M NaCl, and 1 mM DTT. The functionality of RapA^650^ compared to H6-RapA was verified through single-molecule washout experiments (Fig. S1).

E. coli core RNAP (αββ′ω) with a SNAP tag on the C-terminus of β’ (RNAP-SNAP) (kind gift from Robert Landick and Rachel Mooney) was labelled with DY-549 dye as described (10). σ^70^RNAP^549^ holoenzyme used in CoSMoS transcription experiments was prepared by incubating 1.33 µM σ^70^ and 240 nM core RNAP in reconstitution buffer (10 mM Tris-Cl, pH 8.0, 30% glycerol, 0.1 mM EDTA, 100 mM NaCl, 20 mM KCl, 20 μM ZnCl2, 3 mM MgCl2, and 0.6 mM DTT) at 37 °C for 30 min and stored at −20 °C for up to 1 h before use.

### CoSMoS transcription experiments

For the transcription experiments (Fig. 1), single-molecule total internal reflection fluorescence microscopy was performed at excitation wavelengths 488, 532 and 633 nm, for observation of the probe^488^, RNAP^549^, and initial DNA^Cy5^ locations before photobleaching and RapA^650^, respectively. Laser powers and acquisition protocols for each experiment are detailed in Tables S4 and S5. Microscope focus was automatically maintained (53). The temperature of the reaction chamber was maintained at 33.1 ± 0.5 °C using a custom temperature-control system. Transcription reactions were conducted as described (11): in short, single-molecule observations were performed in glass flow chambers (volume ∼20 µL) passivated with succinimidyl (NHS) polyethylene glycol (PEG) and NHS-PEG-biotin (Laysan Bio Inc.; Arab, AL). Streptavidin (#21125; Life Technologies; Grand Island, NY) was introduced at 220 nM in wash buffer (50 mM Tris acetate, 100 mM potassium acetate, 8 mM magnesium acetate, 27 mM ammonium acetate, 0.1 mg mL^−1^ bovine serum albumin (BSA) (#126615 EMB Chemicals; La Jolla, CA), pH 8.0), incubated 1 min, and washed out (all wash-out steps used two flushes each of four chamber volumes of wash buffer). Streptavidin-coated fluorescent beads (T-10711, Molecular Probes), used as markers for stage drift correction, were loaded in the chamber at a dilution of ∼1:400,000 and excess beads were washed out (29). The chamber was then incubated with 25 pM DNA^Cy5^ in wash buffer for 5 min and excess was washed out with buffer. Locations of surface-tethered DNA^Cy5^ molecules were recorded by acquiring five 1 s exposure images with 633 nm excitation at a power of 400 µW (all laser powers measured incident to the objective lens) (53). To photobleach the tethered DNA^Cy5^ molecules, we increased the 633 nm laser power to 1.8 mW and watched the bleaching of DNA^Cy5^ spots for 25-30 minutes. Photobleaching of the DNA molecules allowed for tracking of RapA^650^ molecules during the subsequent transcription reactions. Next, σ^70^RNAP^549^ holoenzyme was introduced into the chamber at 1 nM in transcription buffer (wash buffer supplemented with 3.5% w/v PEG 8,000 (#81268; Sigma-Aldrich; St. Louis, MO), 1 mg mL^−1^ BSA, and an O2-scavenging system, incubated for 10 min to wait for promoter binding and formation of open complexes, and excess unbound RNAP^549^ was washed out. Transcription was initiated by introducing a solution containing 500 µM each of ATP, CTP, GTP, and UTP, 10 nM probe^488^, with or without 5 nM RapA^650^ in transcription buffer. Image acquisition began within 10 s after loading the transcription reagents (at *t* = 0), with excitation alternating between simultaneous 532/633 nm (400 µW each) and 488 nm (400 µW) with a frame rate of 1 frame/s for 40 min.

### CoSMoS rPTC experiments

Microscopy for the experiments in Figs. 2, 3, and 4 was done the same as for the transcription experiments except that we used excitation wavelengths 532 nm to monitor RNAP^549^ and 633 nm to find DNA^Cy5^ locations initially before photobleaching and (in Figs. 3 and 4) afterwards to monitor RapA^650^ (32). For the pre-mix experiments in Fig 2C (orange data), 0.7 nM RNAP^549^, varying concentrations of RapA, and zero or 1 mM ATP were pre-mixed in microcentrifuge tubes and the mixture then introduced into the chamber in transcription buffer at time zero. For the washout experiments in Figs. 2C (green), 3, and 4, 1 nM RNAP^549^ was introduced into the chamber in transcription buffer, incubated for 10 min to wait for binding to npDNA, and excess unbound RNAP^549^ was washed out. Then, varying concentrations of RapA^650^ and 1 mM ATP, 1 mM AMPPNP, or no nucleotide were mixed in transcription buffer and loaded into the glass chamber at time zero. Image acquisition began at time zero for both pre-mix and washout experiments.

### Single-molecule dwell time analyses

Image analysis was done using custom software and algorithms for automatic spot detection, spatial drift correction and co-localization (33). Lifetimes τ in Figs. 1E, 5A, and S4 were fit using the maximum likelihood method to the probability density models

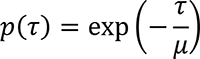

(for Fig. 1E), and to

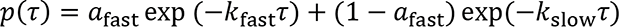

where *k*fast > *k*slow (for Fig. 5A and Fig. S4). Parameter confidence intervals were calculated by bootstrapping and error propagation as described (33).

Characteristic lifetimes of RNAP^549^ in Fig. 2 were calculated by taking the average <τ> of all measured intervals where RNAP^549^ is present on DNA. Intervals that are censored by the maximum acquisition time were included but not corrected in the computation of the average lifetime, as follows:

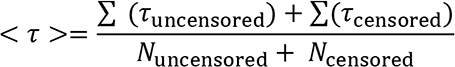

And the standard error of < *τ* >^−1^ was calculated as < *τ* >^−2^ s. e. m. (< *τ* >). Photobleaching rates in the experiment of Fig. 2 (see Table S4) are too low to significantly affect the mean lifetimes.

To measure the rate of RapA^650^ association to either DNA^Cy5^ or RNAP^549^-DNA^Cy5^ complexes (Figs. 3B and 4B), we measured the time of first appearance of RapA^650^ at each DNA^Cy5^ location, RNAP^549^-DNA location (i.e., DNA^Cy5^ locations where RNAP^549^ was already bound), or control no-DNA locations. These data were then fit as described(33) to determine the rate constant *k*ns for the nonspecific association of RapA^650^, the active fraction *A*f of DNA^Cy5^ locations capable of RapA^650^ binding, and the rate constant *k*on,app for association of RapA^650^ with the active fraction of RNAP^549^-DNA sites.

### Simulation of the kinetic model for RNAP dissociation from the PTC by RapA

To analyze our model for dissociation of the RapA•RNAP•DNA complexes based on the Figure S5 reaction scheme, we first defined the row-vector **P**(*t*) to represent the state of the system at time *t*, with each element of the vector representing the probability of the system being in a state as shown here:

| | $P_1(t)$ | $P_2(t)$ | $P_3(t)$ | $P_4(t)$ | $P_5(t)$ |
| --- | --- | --- | --- | --- | --- |
| Molecular species | RNAP•DNA<br>or<br>RapA*•RNAP•DNA | RapA•RNAP•DNA | RapA•RNAP•DNA<sup>†</sup> | DNA +<br>RapA•RNAP | RapA•RNAP*•DNA |
| Initial state | 0 | 1 | 0 | 0 | 0 |

The asterisks represent photobleached components. The time evolution of **P**(*t*) is described by the differential equation where **Q** is the transition rate matrix (54)

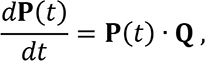

where **Q** is the transition rate matrix (54)

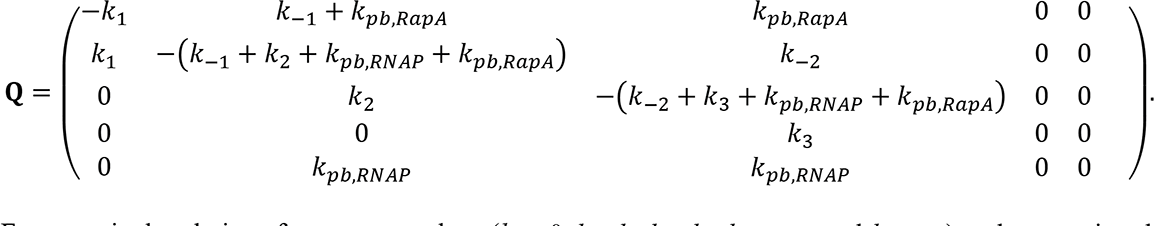

For a particular choice of parameter values (*k*1 = 0, *k*-1, *k*2, *k*-2, *k*3, *k*pb,RNAP, and *k*pb,RapA) and expressing the time *t* = 0 initial state as occupying the molecular species 2 ternary complex (i.e., **P**(0) = (0, 1, 0, 0, 0)), we numerically solved the above differential equation. This yields exact expressions for the time dependence of the **P**(*t*) functions. These **P**(*t*) fractional occupancies are then computed for each molecular species at integral time values *t* ranging from 0 to 3,000 s. We numerically differentiate *P*_1_(*t*), *P*_4_(*t*), and *P*_5_(*t*) to obtain the time-dependent probability densities for molecular species 1, 4, and 5. The three probability densities and the experimental data were then used to compute the likelihood, which was maximized by adjusting the parameters, yielding the rate constant values *k*-1, *k*2, *k*-2, *k*3, *k*ph,RNAP, and *k*ph,RapA given in Fig. 5C. Parameter standard errors were computed by bootstrapping (10,000 samples).

## ACKNOWLEDGEMENTS

We thank Katsuhiko Murakami for generously providing us with the H6-RapA expression plasmid and the RapA protein used in preliminary experiments, Robert Landick and Rachel Mooney for the gift of the RNAP-SNAP protein, and Luke Lavis for the JFX-650 dye adduct. We thank James Portman and Katsuhiko Murakami for comments on the manuscript, and members of the Landick, Hoskins, Campbell, Darst, J.K, and J.G labs for insightful discussions. This work was funded by grants from NIGMS (R01 GM081648 to J.G.) and the Simons Foundation (to J.K).

## Supplementary Information

### SUPPLEMENTARY FIGURES

**Figure S1.**
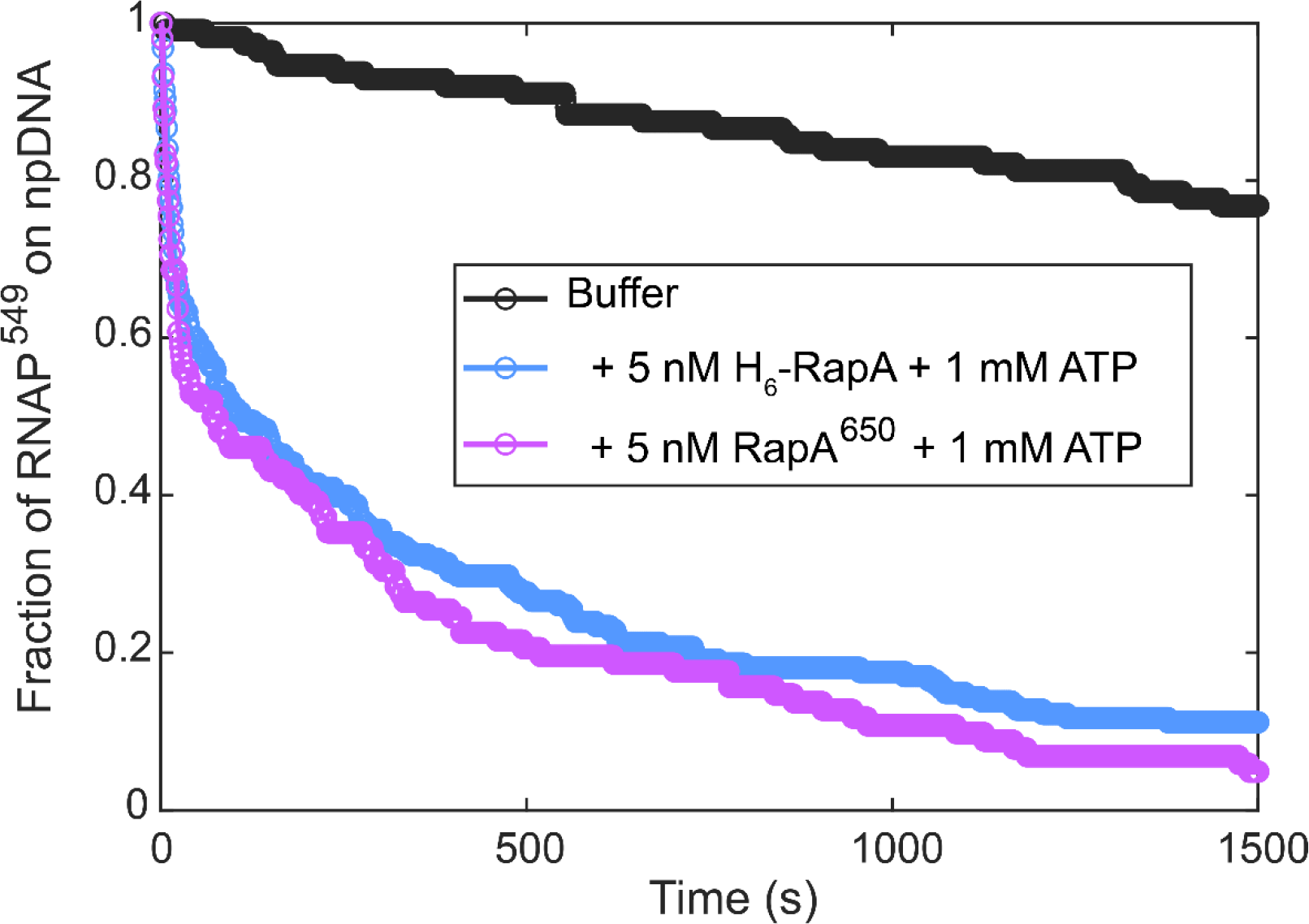
**Single-molecule experiment comparing the activity of labeled RapA^650^ and WT H6-RapA on rPTCs**. Plots show cumulative dwell time distributions of RNAP^549^ in wash buffer in the absence or presence of the indicated RapA constructs.

**Figure S2.**
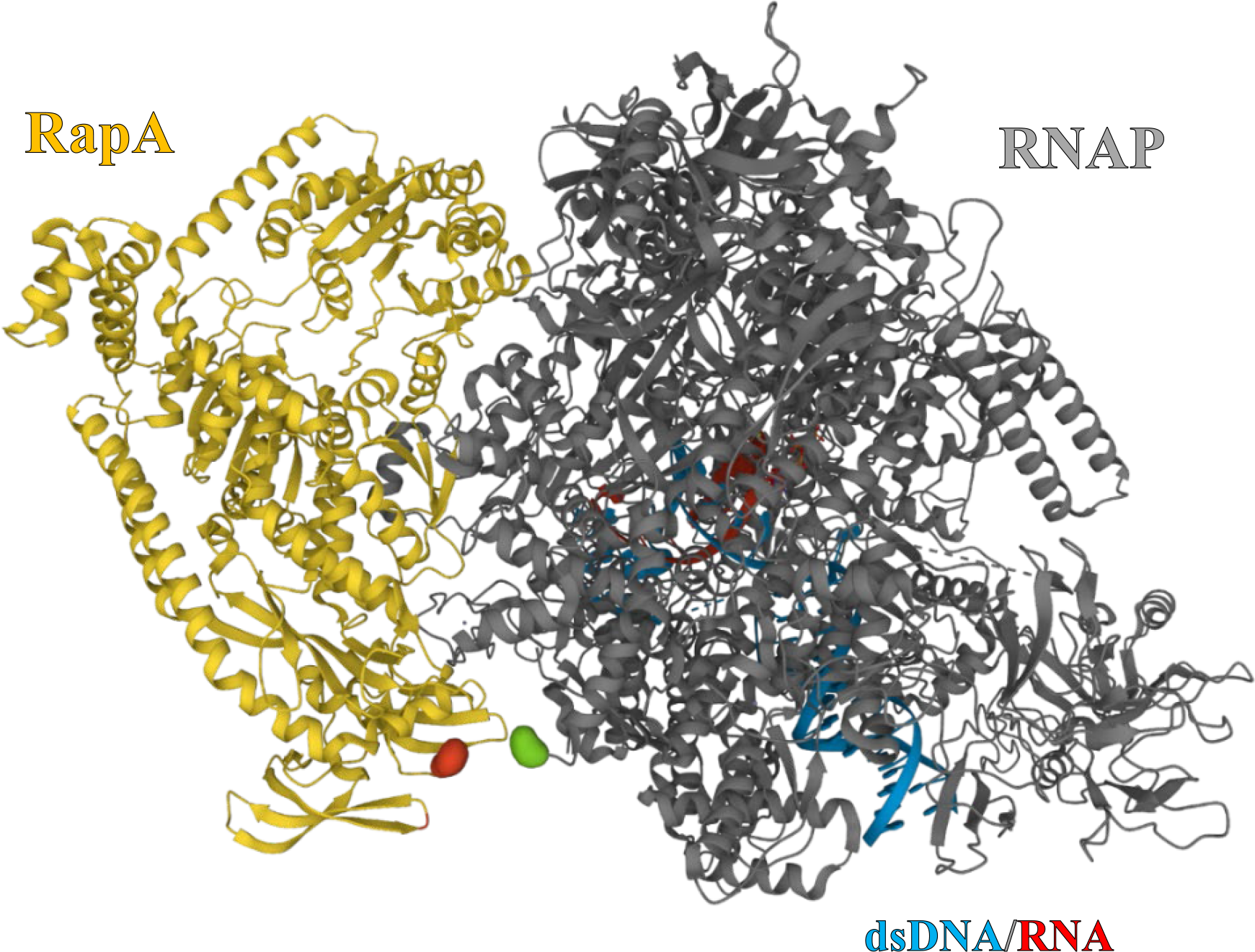
Cryo-EM structural model of RapA complexed to a reconstituted *E. coli* transcription elongation complex. Data are from PDB 7MKN(27); a similar structure is reported in ref. (38) (PDB 7M8E). Illustration highlights the locations of the C-terminus of the RNAP β′ subunit (green) and the N- terminus of RapA (red), which are the attachment points for the SNAP tags in RNAP^549^ and RapA^650^; the termini are 3.3 nm apart. Given the uncertainty about the orientations of the two SNAP tags as well as possible structural differences between the RapA-elongation complex and RapA-rPTC complex we studied, the measured <*E*FRET> = ∼0.4 (Fig. 3D) is plausible given the FRET *R*0 = 4.2 nm for this dye pair.

**Figure S3.**
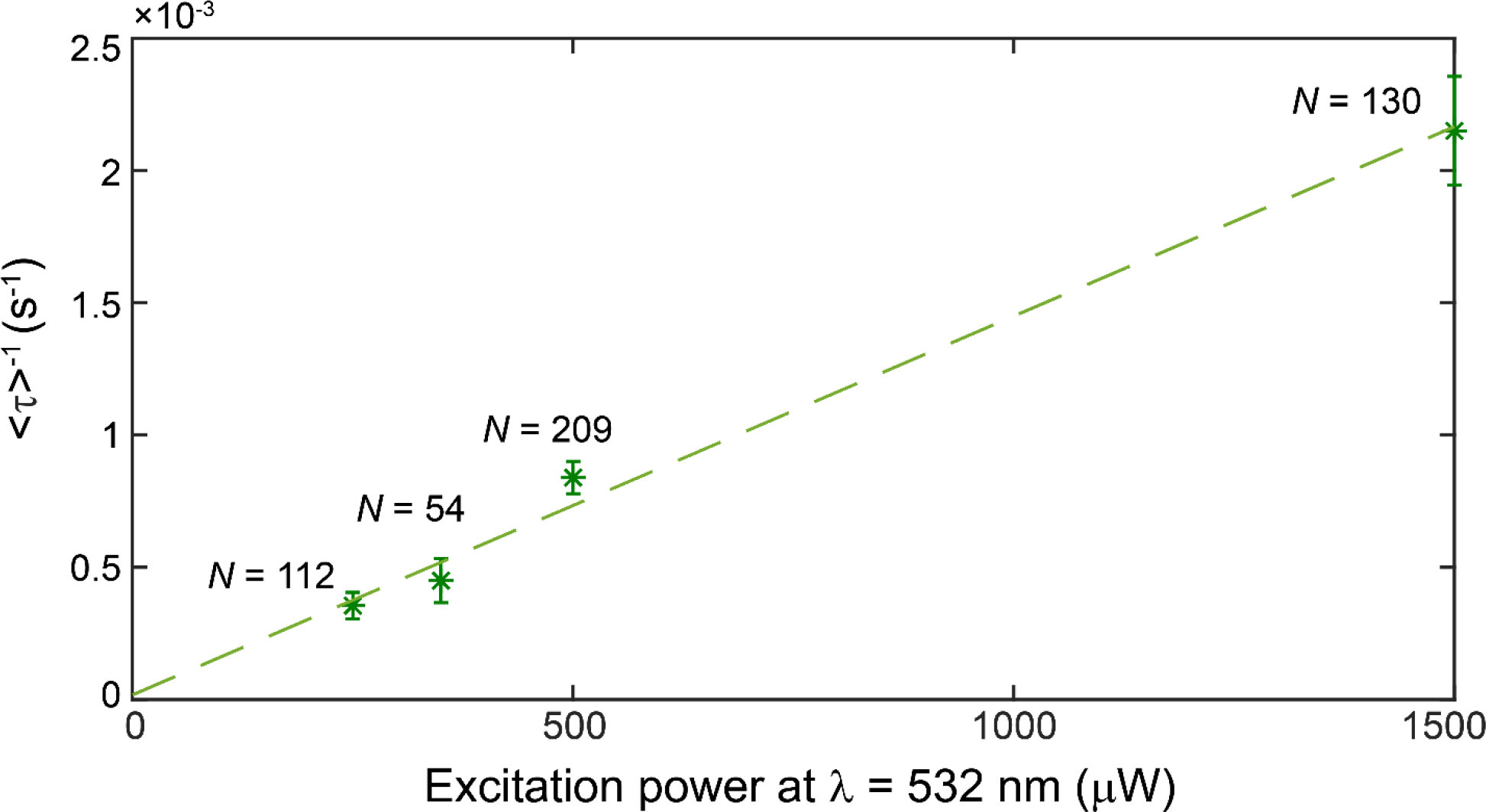
Rate of RNAP^549^ photobleaching. Reciprocal of the mean RNAP^549^ dwell time <τ> in washout experiments (as in Fig. 2) in the absence of RapA, conducted with continuous excitation at different 532 nm laser excitation powers. The linear fit (green line) slope yields the photobleaching-specific rate constant *m*_pb,RNAP_ = 1.43 ± 0.09 × 10^-6^ µW^-1^ s^-1^. RNAP549 photobleaching rates calculated from these results are given in Table S4.

**Figure S4.**
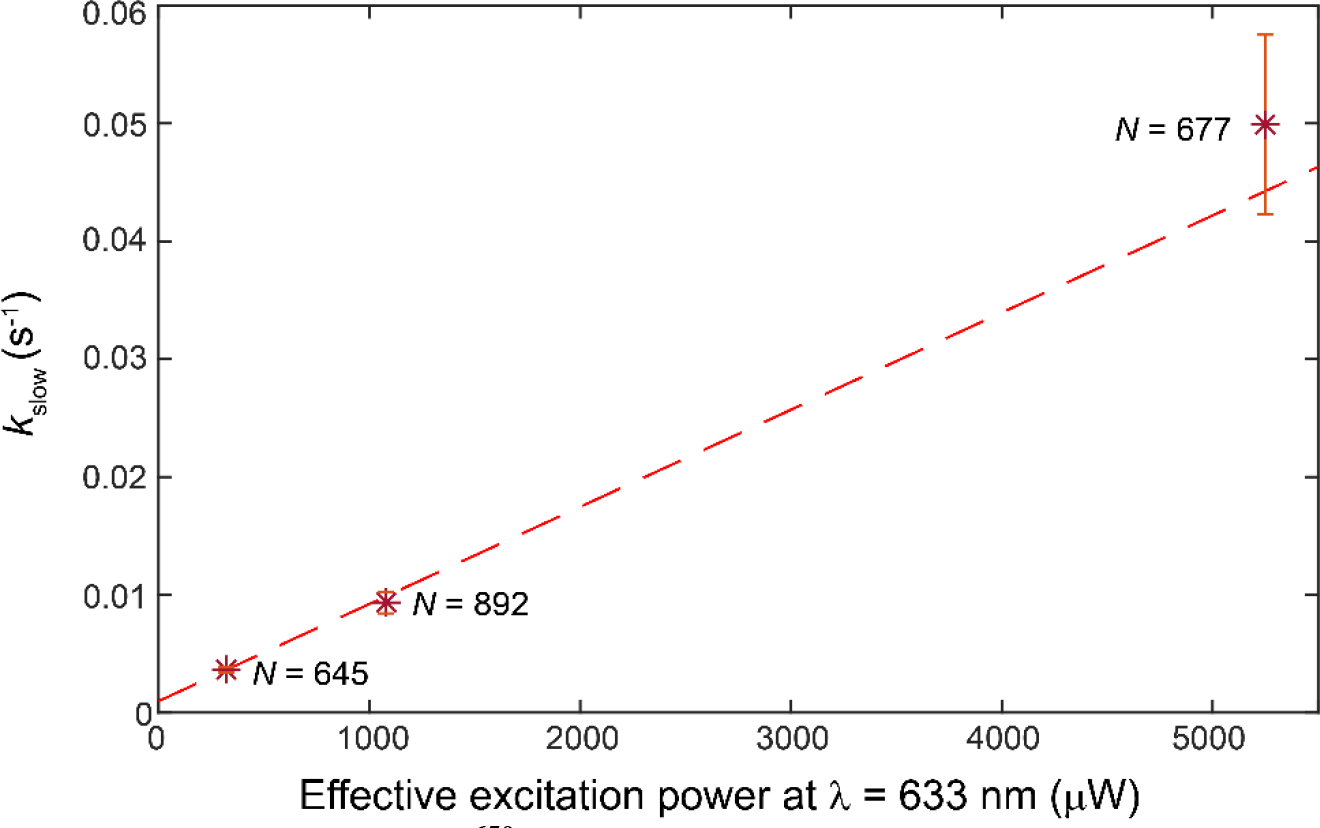
Rate of RapA^650^ photobleaching. The apparent departure rate *k*slow of the long component of the RapA^650^ lifetime distribution in RapA^650^•RNAP^549^•DNA complexes. Data are from the 1 mM AMP- PNP experiment of Fig. 4 and in analogous experiments performed at different 633 nm laser powers. The effective excitation power *P*eff for each experiment is calculated in Table S5. The linear fit slope yields the photobleaching specific rate constant *m*_pb,RapA_ = 8 ± 3 × 10^-6^ µW^-1^ s^-1^, which was used in the calculation of the RapA^650^ photobleaching rate (*k′*pb,RapA) for different experiments (Table S5).

**Figure S5.**
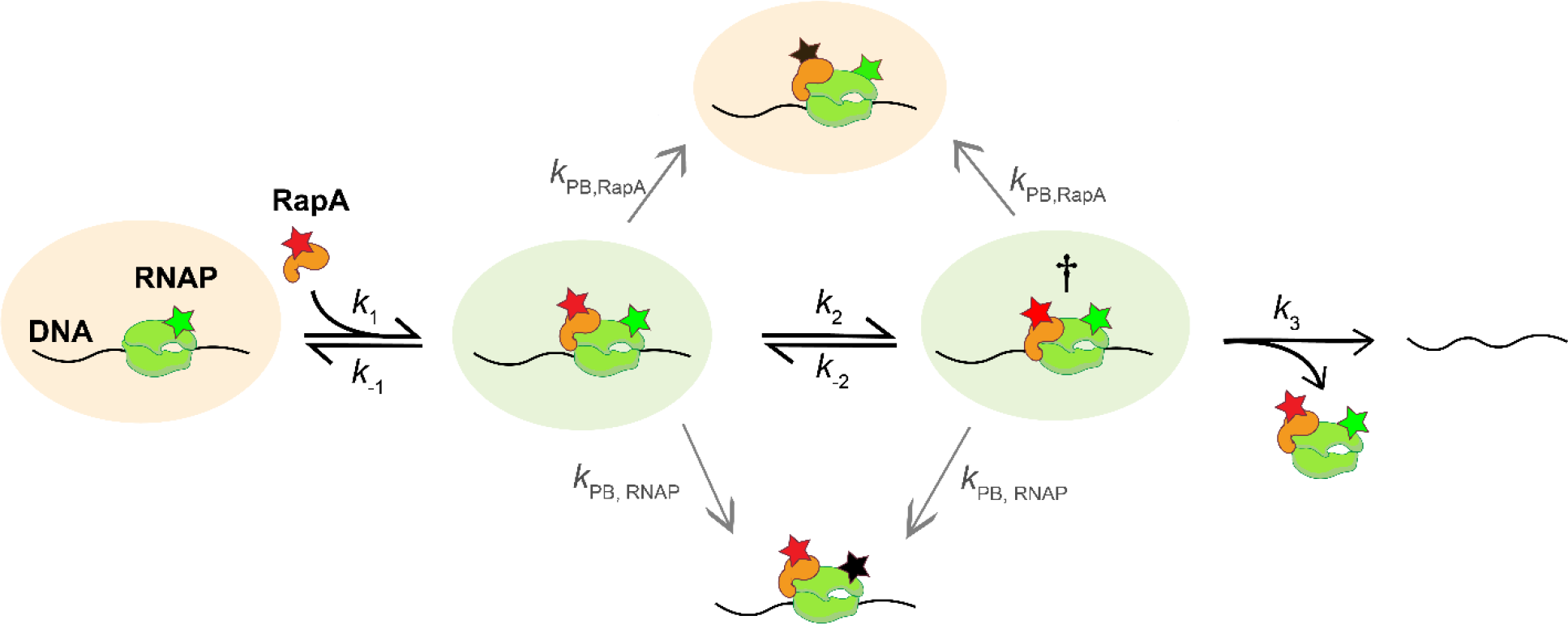
Complete kinetic scheme used for quantitative modeling of the mechanism shown in. Fig. 5B. RNAP (green) and RapA (orange) are illustrated colocalizing with DNA (black). The figure shown here differs from Fig. 5B in that it includes the dye molecules (stars) and explicitly considers RNAP^549^ and RapA^650^ photobleaching, which occur at rates *k*pb,RNAP and *k*pb,RapA, respectively, and yield non-fluorescent molecules (proteins with black stars). This complete kinetic scheme was used to fit the experimental data from Fig. 4 and obtain the rate constants in Fig 5C. Background colors indicate pairs of species that are not distinguishable by fluorescence but whose presence is inferred from the kinetic analysis.

**Figure S6.**
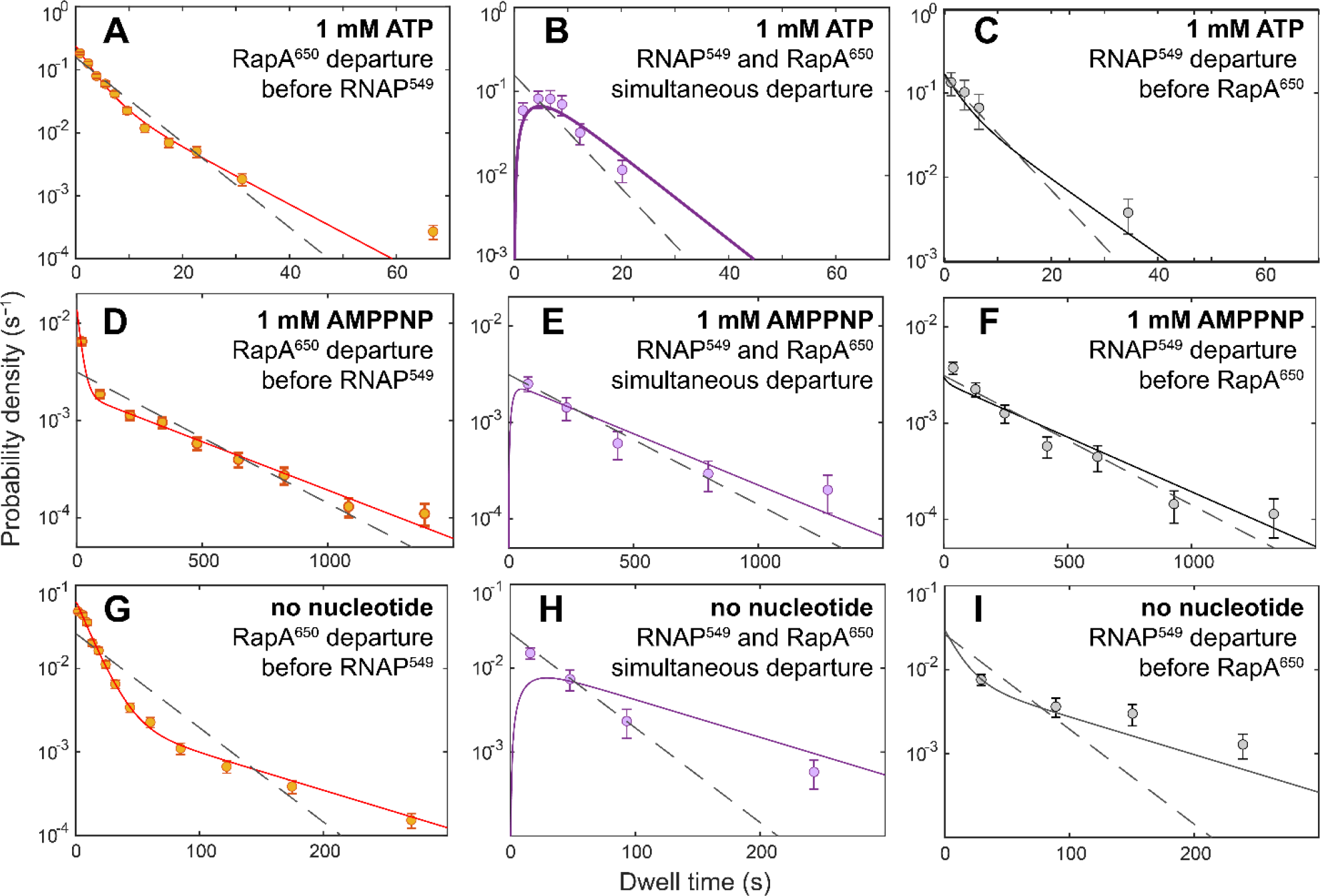
Dwell time distributions of RapA•RNAP•DNA complexes grouped by their fates. RapA•RNAP•DNA dwell time distributions (± S.E.; circles) were measured in the three different nucleotide condition experiments in Fig 4: 1 mM ATP **(A, B, C)**, 1 mM AMPPNP **(D, E, F)**, and no added nucleotide **(G, H, I)**. For each experiment, measured dwell times were separated into three groups according to the fates of the RapA•RNAP•DNA complexes: RapA^650^ departs first **(A, D, G),** RNAP^549^ and RapA^650^ depart simultaneously **(B, E, H)**, and (3) RNAP^549^ departs first **(C, F, I)**. Number of observed dwell times in each distribution are reported in Table S6. Also plotted are fits to the kinetic scheme including photobleaching processes (Fig. S5), which are consistent (solid lines) with the data. Fits (dashed lines) to scheme a with only one species of RapA•RNAP•DNA ternary complex (Fig. S7) are inconsistent with the data.

**Figure S7.**
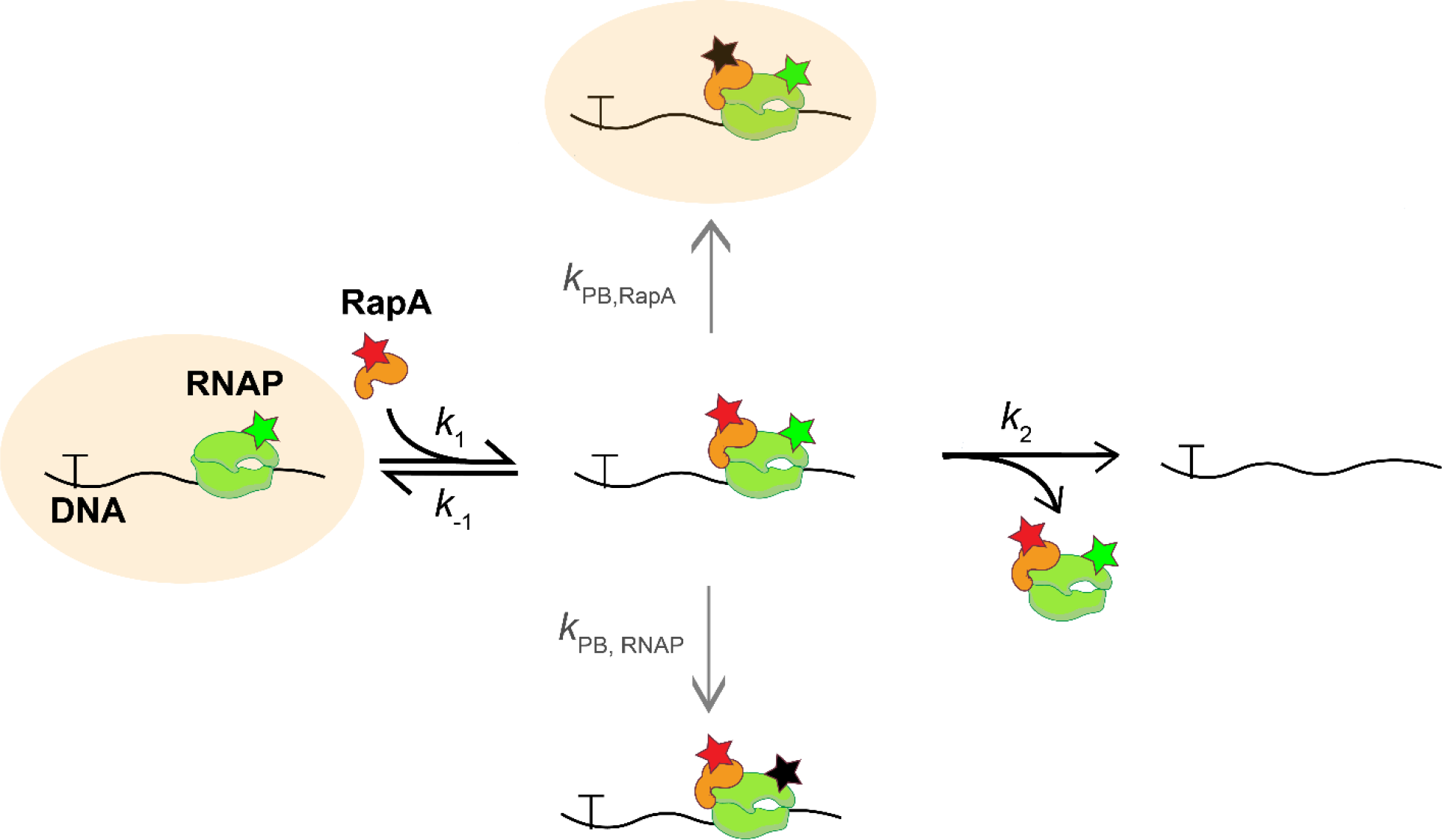
Kinetic scheme with a single ternary complex. A complete model including photobleaching (illustrated as in Fig. S5) but with only one ternary RapA•RNAP•DNA complex. Fits to this scheme (Fig S6, dashed lines) but are inconsistent with the data (Fig. S6, circles).

### SUPPLEMENTARY TABLES

**Table S1:** RNAP^549^ dwell time on npDNA as a function of RapA^650^ concentration (**Fig. 2C**)

| Condition | $k_{\max}$<br>($\times 10^{-3} \text{ s}^{-1}$) | $K_{0.5}$<br>(nM) | $b$<br>($\times 10^{-3} \text{ s}^{-1}$) | $m$<br>($\times 10^6 \text{ M}^{-1} \text{ s}^{-1}$) |
| --- | --- | --- | --- | --- |
| Pre-mix + ATP* | $43 \pm 4$ | $21 \pm 3$ | $1.8 \pm 0.1$ | -- |
| Washout + ATP<sup>†</sup> | -- | -- | $0.78 \pm 0.02$ | $1.4 \pm 0.07$ |
| Premix; no ATP<sup>†</sup> | -- | -- | $1.84 \pm 0.09$ | $0.14 \pm 0.03$ |
| Washout; no ATP<sup>†</sup> | -- | -- | $0.62 \pm 0.08$ | $0.002 \pm 0.002$ |

\*Data fit to a hyperbolic model of the form $\langle \tau \rangle^{-1} = b + \frac{k_{\max}[\text{RapA}]}{K_{0.5} + [\text{RapA}]}$ .
<sup>†</sup> Data fit to a linear model of the form $\langle \tau \rangle^{-1} = m[\text{RapA}] + b$ .

**Table S2:** Kinetics of RapA^650^ binding at sites with rPTC or npDNA*

| Binding Target | $A_f$ | $k_{\text{on,app}}$<br>($\times 10^{-3} \text{ s}^{-1}$) | $k_{\text{on}}$<br>($\times 10^6 \text{ s}^{-1} \text{ M}^{-1}$) | $k_{\text{ns}}$<br>($\times 10^{-3} \text{ s}^{-1}$) | $N_{\text{DNA}}$ | $N_{\text{ns}}$ |
| --- | --- | --- | --- | --- | --- | --- |
| rPTC | $0.73 \pm 0.05$ | $9 \pm 2$ | $1.8 \pm 0.4$ | $0.40 \pm 0.02$ | 141 | 327 |
| npDNA | $0.02 \pm 0.01$ | $98 \pm 180^{**}$ | $20 \pm 36^{**}$ | $0.29 \pm 0.03$ | 42 | 85 |

\*See Fig. 3B. $A_f$ , inferred active fraction of target molecules capable of specific RapA binding; $k_{\text{on,app}}$ , apparent first-order rate constant for specific binding of RapA at rPTC or npDNA locations; $k_{\text{on}}$ , second-order binding rate constant calculated by dividing $k_{\text{on,app}}$ by the concentration of RapA<sup>650</sup> in the sample; $k_{\text{ns}}$ , apparent first-order rate constant for non-specific binding to the chamber surface; $N_{\text{DNA}}$ , number of target locations; $N_{\text{ns}}$ , number of off-target locations.
\*\*Not distinguishable from zero.

**Table S3:** Kinetics for FRET-detected RapA^650^ association with rPTC*

| **Table S3: Kinetics for FRET-detected RapA<sup>650</sup> association with rPTC\*** |  |  |  |  |  |  |
| --- | --- | --- | --- | --- | --- | --- |
| Condition | $A_f$ | $k_{on,app}$<br>($\times 10^{-3} s^{-1}$) | $k_1^{**}$<br>($\times 10^6 s^{-1} M^{-1}$) | $k_{ns}$<br>($\times 10^{-5} s^{-1}$) | $N_{DNA}$ | $N_{ns}$ |
| 1 mM ATP | $0.71 \pm 0.03$ | $8 \pm 1$ | $1.6 \pm 0.2$ | $6.0 \pm 0.8$ | 289 | 64 |
| 1 mM AMP-PNP | $0.80 \pm 0.02$ | $9.1 \pm 0.7$ | $1.8 \pm 0.2$ | $9 \pm 1$ | 336 | 201 |
| no nucleotide | $0.71 \pm 0.03$ | $8.0 \pm 0.9$ | $1.6 \pm 0.2$ | $6.9 \pm 0.8$ | 309 | 208 |

\*Fits of data plotted in Fig. 4B. $A_f$ , inferred active fraction of rPTCs capable of specific RapA association; $k_{on,app}$ , apparent first-order rate constant for specific binding of RapA at rPTC or npDNA locations; $k_{on}$ , second-order binding rate constant calculated by dividing $k_{on,app}$ by the concentration of RapA<sup>650</sup> in the sample; $k_{ns}$ , apparent first-order rate constant for non-specific binding to the chamber surface; $N_{DNA}$ , number of target DNA molecule locations; $N_{ns}$ , number of control off-DNA locations.

**Table S4:** RNAP^549^ photobleaching rates calculated for different experimental conditions

|  |  |  |  |  |
| --- | --- | --- | --- | --- |
| Experiment | 533 nm laser<br>excitation<br>power, $I$ ($\mu W$) | Laser duty<br>ratio, $D^*$ | Effective exposure, $E_{eff}$<br>($\mu W$)<sup>†</sup> | Photobleaching rate,<br>$k'_{pb,RNAP}$<br>($\times 10^{-4} s^{-1}$)<sup>‡</sup> |
| Fig. 1 | 400 | 0.42 | 167 | $2.4 \pm 0.2$ |
| Fig. 2 | 400 | 1.0 | 400 | $5.7 \pm 0.4$ |
| Fig. 3A,B | 2,100 | 0.2 | 420 | $6.0 \pm 0.4$ |
| Fig. 3C,D | 2,100 | 0.2 | 420 | $6.0 \pm 0.4$ |
| Fig. 4 | 2,100 | 0.14 | 300 | $4.3 \pm 0.3$ |

\*Fraction of time the sample is exposed to the 532 nm laser. <sup>†</sup> $E_{eff} = I \times D$ . <sup>‡</sup> $k'_{pb,RNAP} = E_{eff} m_{pb,RNAP}$ , where $m_{pb,RNAP}$ is the photobleaching specific rate constant, $1.43 \pm 0.09 \times 10^{-6} \mu W^{-1} s^{-1}$ (Fig. S3). This calculation overestimates $k'_{pb,RNAP}$ in experiments where FRET to RapA<sup>650</sup> partially quenches RNAP<sup>549</sup>.

**Table S5:** RapA^650^ photobleaching rates calculated for different experimental conditions

| | Laser<br>excitation<br>power, $P_{633}$<br>($\mu\text{W}$) | Laser<br>excitation<br>power, $P_{532}$<br>($\mu\text{W}$) | Duty ratio,<br>$D^*$ | Effective 633 nm<br>exposure, $P_{\text{eff}}$<br>($\mu\text{W}$)<sup>†</sup> | Photobleaching<br>rate, $k'_{\text{pb,RapA}}$<br>($\times 10^{-3} \text{ s}^{-1}$)<sup>‡</sup> |
| --- | --- | --- | --- | --- | --- |
| Figure 1 | 400 | 400 | 0.42 | 250.5 | $2.1 \pm 0.7$ |
| Figure 3A,B | 1,500 | 1,500 | 1 | 2,250 | $18 \pm 6$ |
| Figure 4;<br>Figure S4<br>(lowest power) | 1,500 | 1,500 | 0.14 | 321 | $2.6 \pm 0.9$ |
| Figure S4<br>(middle power) | 0 | 2,160 | 1 | 1,080 | $8.9 \pm 2.9$ |
| Figure S4<br>(highest power) | 3,500 | 3,500 | 1 | 5,250 | $40 \pm 10$ |

\*Fraction of time the sample is exposed to the 633 or 532 nm laser; $D_{532} = D_{633} = D$ .
<sup>†</sup> $P_{\text{eff}} = P_{633}D_{633} + P_{532}D_{532} \frac{\text{Em}_A\text{Ex}_{532}}{\text{Em}_A\text{Ex}_{633}}$ , where $\text{Em}_A\text{Ex}_{532}$ is the emission intensity in the acceptor channel from exciting a donor/acceptor pair at 532 nm, and $\text{Em}_A\text{Ex}_{633}$ is the emission intensity in the acceptor channel from exciting a donor/acceptor pair at 633 nm.
<sup>‡</sup> $k'_{\text{pb,RapA}} = P_{\text{eff}} m_{\text{pb,RspA}}$ , where $m_{\text{pb,RspA}}$ is the photobleaching specific rate constant, $8 \pm 3 \times 10^{-6} \mu\text{W}^{-1} \text{ s}^{-1}$ (Fig. S4).

**Table S6:** Fates of RNAP^549^·RapA^650^·DNA complexes measured from data (**Fig. 4**) and predicted by the fit to the complete kinetic scheme (Fig. S5)*

| Nucleotide condition | Number of complexes | Number of observations (percent of total ± S.E.) | | |
| --- | --- | --- | --- | --- |
|  |  | Percent of total calculated from fit |  |  |
|  |  | RapA<sup>650</sup> departure before RNAP<sup>549</sup> (Fate #1) | RNAP<sup>549</sup> and RapA<sup>650</sup> simultaneous departure (Fate #2) | RNAP<sup>549</sup> departure before RapA<sup>650</sup> (Fate #3)<sup>†</sup> |
| 1 mM ATP | 647 | 513 (79 ± 2%)<br>85 ± 1% | 114 (18 ± 2%)<br>15 ± 1% | 20 (2 ± 1%)<br>0.30 ± 0.03% |
| 1 mM AMPPNP | 645 | 446 (69 ± 2%)<br>73 ± 2% | 85 (13 ± 1%)<br>11 ± 1% | 115 (18 ± 2%)<br>15 ± 1% |
| None | 1,305 | 1,208 (93 ± 1%)<br>98.0 ± 0.2% | 43 (3 ± 1%)<br>0.14 ± 0.01% | 54 (4 ± 1%)<br>1.4 ± 0.1% |

\*Entries with white background are experimental results; entries with grey background are calculated from the fit.
<sup>†</sup>This fate is completely accounted for by RNAP<sup>549</sup> photobleaching (see text and Fig 5C) .

**Table S7:** Fit parameters for RapA•RNAP•DNA lifetime distributions (Fig. 5A)*

| Nucleotide condition | *N* | Amplitude, *a*<sub>fast</sub> | *k*<sub>fast</sub> (s<sup>-1</sup>) | *k*<sub>slow</sub> (s<sup>-1</sup>) |
| --- | --- | --- | --- | --- |
| 1 mM ATP | 647 | 0.76 ± 0.04 | 0.228 ± 0.009 | 0.029 ± 0.005 |
| 1 mM AMPPNP | 645 | 0.35 ± 0.03 | 0.057 ± 0.007 | 0.0036 ± 0.0002 |
| None | 1,305 | 0.81 ± 0.03 | 0.049 ± 0.003 | 0.0071 ± 0.0009 |

\*Parameters ( ± S.E.) are from fits to a two-exponential model (see Materials and Methods).

## Notes

### Competing Interest Statement

The authors have declared no competing interest.

